# Cerebrospinal fluid YWHAG:NPTX2 ratio predicts clinical severity and future phenoconversion in frontotemporal lobar degeneration

**DOI:** 10.64898/2026.08.22.746470

**Authors:** Hamilton Se-Hwee Oh, Joshua D. Downer, Connor D. Dietz, Claire Yballa, Edoardo Marcora, Argentina Lario-Lago, Hilary W. Heuer, Leah K. Forsberg, Jennifer Hsiao-Nakamoto, Chi-Lu Chiu, Paul Auger, Casey Powers, Gil Di Paolo, Fen Huang, Brian Appleby, Sami Barmada, Ece Bayram, Andrea Bozoki, David Clark, R. Ryan Darby, Bradford Dickerson, Kimiko Domoto-Reilly, Kelley Faber, Anne Fagan, Tatiana Foroud, Douglas R. Galasko, Daniel Geschwind, Nupur Ghoshal, Neill Graff-Radford, Ian M. Grant, Chadwick M Hales, Lawrence S. Honig, Ging-Yuek Hsiung, Edward D. Huey, David Irwin, David Knopman, John Kornak, Justin Kwan, Gabriel C. Léger, Irene Litvan, Ian R. Mackenzie, Mario F. Mendez, Chiadi Onyike, Belen Pascual, Peter Pressman, Rosa Rademakers, Eliana Marisa Ramos, Erik D. Roberson, Allison Snyder, Carmela Tartaglia, Dylan Wint, Arabella Bouzigues, Lucy L. Russell, Phoebe H. Foster, Eve Ferry-Bolder, John C. van Swieten, Lize C. Jiskoot, Harro Seelaar, Raquel Sanchez-Valle, Robert Laforce, Caroline Graff, Daniela Galimberti, Rik Vanderberghe, Alexandre de Mendonça, Giuseppe di Fede, Isabel Santana, Alexander Gerhard, Johannes Levin, Benedetta Nacmias, Markus Otto, Maxime Bertoux, Thibaud Lebouvier, Simon Ducharme, Chris R. Butler, Isabelle Le Ber, Elizabeth Finger, Mario Masellis, James B. Rowe, Matthis Synofzik, Fermin Moreno, Barbara Borroni, Aitana Sogorb-Esteve, Sophia Weiner, Johan Gobom, Henrik Zetterberg, Jonathan D. Rohrer, Emily W. Paolillo, Lawren VandeVrede, Peter J. Ljubenkov, Renaud La Joie, Gil D. Rabinovici, Joel H. Kramer, Bruce L. Miller, Maria Luisa Gorno-Tempini, Tania Gendron, Leonard Petrucelli, Bradley F. Boeve, Howard J. Rosen, Jennifer S. Yokoyama, Scott J. Russo, William W. Seeley, Julio C. Rojas, Kaitlin B. Casaletto, Adam M. Staffaroni, Alison Goate, Adam L. Boxer, Rowan Saloner, the Frontotemporal Dementia Prevention Initiative (FPI)

## Abstract

Frontotemporal lobar degeneration (FTLD) is a common cause of early-onset dementias marked by progressive declines in behavior, cognition, and/or movement. FTLD neuropathologies, including TDP-43 proteinopathies and primary tauopathies, do not have reliable fluid biomarkers for in-vivo diagnosis nor biomarkers that directly correspond to FTLD clinical features. Fluid biomarkers that forecast and track FTLD clinical progression, irrespective of pathology or clinical syndrome, are urgently needed to improve clinical trial designs. We previously identified the ratio between two cerebrospinal fluid (CSF) synaptic proteins, YWHAG and NPTX2, as a prognostic biomarker of cognitive decline in Alzheimer’s disease (AD), independent of core AD pathologies, amyloid and tau. Here, we evaluate its utility in sporadic and familial FTLD compared to other neurodegenerative diseases. Using CSF assays from four independent cohorts (UCSF-MAC, ALLFTD, GENFI, PDBP), we find CSF YWHAG:NPTX2 is substantially elevated across all sporadic and familial FTLD syndromes, AD, and dementia with Lewy bodies. CSF YWHAG:NPTX2 robustly correlates with clinical severity across sporadic and familial FTLD (*C9orf72*, *GRN*, or *MAPT* mutations), independent of current gold-standard neurodegeneration biomarker neurofilament light (NfL). In presymptomatic familial FTLD, CSF YWHAG:NPTX2 is estimated to rise roughly a decade before symptom onset and improves prediction of imminent symptomatic conversion by 1.7-fold compared to plasma NfL alone, more than halving the estimated sample size required for an FTLD prevention clinical trial. These findings underscore CSF YWHAG:NPTX2 as a cross-dementia synaptic biomarker of cognitive decline and a promising biomarker for disease staging and prognosis across the clinico-pathological continuum of FTLD.

## MAIN

Frontotemporal lobar degeneration (FTLD) is a common cause of early age-of-onset neurodegenerative disorders, collectively accounting for approximately 10-20% of dementia cases under the age of 65^1,2^. The majority of FTLD is pathologically defined by either accumulation of microtubule binding protein tau (FTLD-tau, ∼45%) or abnormal deposits of transactive response DNA-binding protein 43 (FTLD-TDP, ∼50%)^3^, with a smaller proportion defined by abnormal accumulation of fused-in-sarcoma protein (FTLD-FET, ∼5%)^4^. 20-30% of FTLD is familial, most often caused by autosomal dominant mutations in *C9orf72*, *GRN*, or *MAPT*^3,5^. Clinically, FTLD is highly heterogeneous with patients varying along a spectrum of progressive behavioral, language, and/or motoric features^3,6^. Patients with the same pathology can manifest disparate syndromes, and, conversely, patients with the same syndrome can harbor different pathologies^7^. Rates of clinical progression and brain atrophy are also highly variable^8^, even within phenotypes or genetic subtypes^9^. Antemortem molecular biomarkers that accurately stratify disease stage and predict the onset and rate of symptom progression are urgently needed to define target patient populations for FTLD clinical trials and care^5,10^.

Fluid biomarkers have transformed the study and management of neurodegenerative diseases by offering a real-time window into molecular changes that precede clinical symptoms. In Alzheimer’s disease (AD), cerebrospinal fluid (CSF) and plasma measures of amyloid and tau (e.g., Aβ42/Aβ40, pTau181, pTau217, MTBR-tau243) and neurofilament light (NfL) are used to track disease pathology and neurodegeneration^11,12^. NfL is the most validated and used biomarker for FTLD-related neurodegeneration and disease progression^13,14^. Despite its utility, NfL does not fully capture FTLD clinical severity^15^, underscoring the need for complementary FTLD biomarkers reflecting additional biological processes underlying clinical severity. We recently identified the CSF YWHAG:NPTX2 ratio as a synaptic protein biomarker that robustly predicts AD onset and progression, independent of Aβ, tau, and NfL^16^. In AD, YWHAG:NPTX2 increases 10-20 years before symptom onset and accelerates with symptom progression^16^. Given substantial evidence linking neurodegenerative diseases to synaptic dysfunction and specific evidence linking YWHAG and NPTX2 individually to FTLD symptoms^17–19^, we hypothesized that this ratio would exhibit strong associations with FTLD clinical stage and importantly, improve prediction of symptom onset in presymptomatic FTLD.

Here, we investigate the CSF YWHAG:NPTX2 ratio in sporadic and familial FTLD and other neurodegenerative diseases using aptamer-based and mass spectrometry-based CSF assays in four independent cohorts: the UCSF Fein Memory and Aging Center (MAC), ARTFL-LEFFTDS Longitudinal Frontotemporal Lobar Degeneration (ALLFTD) Consortium, Genetic Frontotemporal Dementia Initiative (GENFI), and Parkinson’s Disease Biomarker Program (PDBP). We demonstrate CSF YWHAG:NPTX2 as a robust marker of clinical severity stage, independent of established biomarkers (e.g., NfL, AD biomarkers), across multiple neurodegenerative conditions, including sporadic and familial FTLD. In presymptomatic familial FTLD, we demonstrate its utility in forecasting imminent conversion to symptomatic disease and continued symptomatic decline compared to NfL alone, thereby highlighting its potential as a predictive and clinical trial stratification biomarker.

## RESULTS

### CSF YWHAG:NPTX2 is markedly elevated across dementias, independent of NfL and core AD biomarkers

We first analyzed data from 384 individuals in the UCSF-MAC cohort to evaluate CSF YWHAG:NPTX2 (SomaScan) differences across sporadic FTLD (n=144), biomarker-confirmed symptomatic AD (n=149), motor neuron disease (MND; n=5) and cognitively unimpaired (CU) controls (n=86; **Fig. 1a, Extended Data Fig. 1a-b, Supplementary Table 1**). CSF YWHAG:NPTX2 levels were markedly increased in FTLD (mean difference (m.d.) = 1.79 standard deviations (s.d.) [1.54–2.03 95% CI], p = 8.8x10^−38^) and AD (m.d. = 2.30 [2.05–2.54] s.d., p = 3.4x10^−55^) versus controls, controlled for age and sex (**Supplementary Table 2**). This was true for all clinical subtypes of FTLD and AD, including amnestic (early-onset AD [EOAD] and late-onset AD [LOAD]), behavioral (i.e., bvFTD, bvAD), primary progressive aphasias (PPA; logopenic [lvPPA], non-fluent [nfvPPA], semantic [svPPA]), movement (corticobasal syndrome [CBS], progressive supranuclear palsy syndromes [PSPs]), and visuocognitive (posterior cortical atrophy [PCA]) presentations (**Fig. 1b, Extended Data Fig. 1c**). Notably, CSF YWHAG:NPTX2 levels were not significantly elevated in the small subset of MND cases without dementia (m.d. = 0.53 [–0.50 to 1.55] s.d., p = 0.20) but elevated in FTD cases with MND (FTD-MND; m.d. = 3.11 [2.86–3.35] s.d., p = 1.4x10^−11^; **Fig. 1b**).

**Figure 1.**
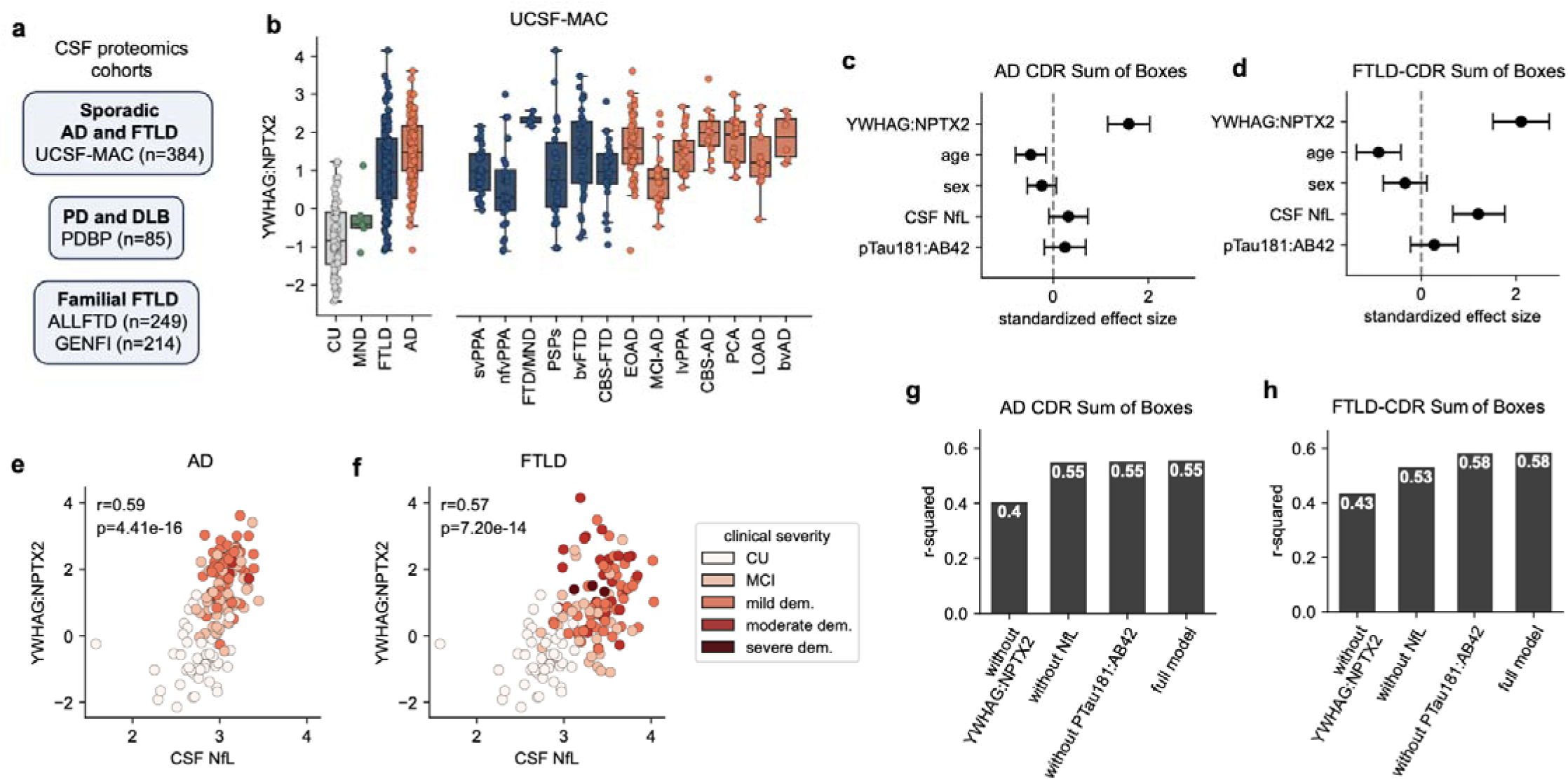
CSF YWHAG:NPTX2 ratio is markedly elevated across sporadic FTLD and AD, independent of NfL. **a**, Study cohorts and sample sizes. **b,** CSF YWHAG:NPTX2 levels across clinical diagnoses including FTLD (n=144) and AD (n=149) spectrum cases, MND (n=5), and cognitively unimpaired controls (n=86) from UCSF-MAC. The box bounds are the Q1, median and Q3; the whiskers show Q1 − 1.5× the interquartile range (IQR) and Q3 + 1.5× the IQR. **c,** Standard CDR Sum of Boxes regressed against CSF YWHAG:NPTX2, age, sex, CSF NfL (Roche Elecsys), and CSF pTau181:AB42 (Roche Elecsys) in a linear model, among AD and control samples (n=154). Points and error bars represent standardized effect sizes and 95% confidence intervals. **d,** As in **c,** but with CDR+NACC-FTLD (FTLD-CDR) Sum of Boxes among FTLD and control samples (n=145). **e,** Linear correlation and p-value between CSF YWHAG:NPTX2 and CSF NfL (log_10_) in AD and control samples, color-coded according to CDR Global (n=154). **f,** As in **e**, but in FTLD and control samples, color-coded according to FTLD-CDR Global (n=145). **g,** R-squared results from a linear model regressing CDR Sum of Boxes against covariates displayed on the x-axis in AD and control samples. Full model includes CSF YWHAG:NPTX2, age, sex, CSF NfL, and CSF pTau181:AB42 (n=154). **h,** as in **g**, but with FTLD-CDR Sum of Boxes in FTLD and control samples (n=145).

We next tested the association between CSF YWHAG:NPTX2 and dementia clinical severity using linear models regressing the Clinical Dementia Rating Sum of Boxes (CDR-SB) against CSF YWHAG:NPTX2, including CSF NfL (Roche Elecsys), CSF pTau181:Aβ42 (Roche Elecsys), age, and sex as covariates in separate models for AD and FTLD. For AD analyses, 154 AD and control subjects with complete values for all covariates were included. For FTLD analyses, 145 FTLD and control subjects with complete values for all covariates were included. For all downstream clinical severity analyses, we used the standard CDR for AD and the CDR®+NACC-FTLD for FTLD, given the CDR®+NACC-FTLD captures additional behavioral and language domains commonly affected in FTLD. CSF YWHAG:NPTX2 showed the strongest association with clinical severity in both AD (standardized β = 1.59 [1.15–2.03], p = 4.7x10^−11^) and FTLD (standardized β = 2.11 [1.52–2.69], p = 6.0x10^−11^), while CSF NfL (standardized β = 1.21 [0.66–1.77], p = 3.0x10^−5^) was independently associated with clinical severity only in FTLD (**Fig. 1c-d**).

Results were similar in sex-stratified (**Extended Data Fig. 2a-b**) and clinical-severity-stratified (**Extended Data Fig. 2c-d**) sensitivity analyses. We also compared CSF YWHAG:NPTX2 with previously investigated CSF proteins (SomaScan measures) representing complementary aspects of neuroinflammation (YKL-40), synaptic function (GAP43), and lysosomal function (PGRN). In joint models with CSF YWHAG:NPTX2, none of these additional CSF proteins showed a significant independent association with clinical severity in either AD or FTLD, while CSF YWHAG:NPTX2 remained the strongest correlate (**Extended Data Fig. 2e**).

Visualization of CSF YWHAG:NPTX2 versus CSF NfL showed moderate correlation in both AD (r=0.59) and FTLD (r=0.57) and confirmed CSF NfL’s independent effect only in FTLD (**Fig. 1e-f**). To compare the added value of CSF YWHAG:NPTX2, CSF NfL, and pTau181:Aβ42 in tracking clinical severity, we compared the variance explained by full and reduced models. In AD, removing CSF YWHAG:NPTX2 decreased variance explained by 15% (55% vs. 40%), while removing CSF NfL or CSF pTau181:Aβ42 did not change variance explained (55% vs. 55%) (**Fig. 1g**). In FTLD, removing CSF YWHAG:NPTX2 decreased variance explained by 15% (58% vs. 43%) and removing CSF NfL by 5% (58% vs. 53%) (**Fig. 1h**). We confirmed that the ratio between YWHAG and NPTX2 provides stronger associations with clinical severity than either protein alone (**Extended Data Fig. 3a-e**). Together, these models highlight CSF YWHAG:NPTX2 as a stronger independent correlate of clinical severity across AD and FTLD.

For AD models, we performed a sensitivity analysis in a subset of 55 AD patients with amyloid and tau positron emission tomography (PET) imaging performed within 6 months of CSF collection (10 mild cognitive impairment [MCI], 2 LOAD, 26 EOAD, 10 PCA, 7 lvPPA). CSF YWHAG:NPTX2 significantly correlated with tau PET temporal meta-ROI values (*r*=0.33, *p*=1.32x10^−2^) but not amyloid PET centiloids (*r*=0.08, *p*=5.86x10^−1^; **Extended Data Fig. 4a-c**). In a linear model regressing CDR-SB against CSF YWHAG:NPTX2, tau PET, amyloid PET, age, and sex, CSF YWHAG:NPTX2 was the only significant predictor (**Extended Data Fig. 4d)**. Removing CSF YWHAG:NPTX2, but not tau PET, dramatically reduced the percent variance in CDR-SB explained by the model (30% vs. 5% after removing CSF YWHAG:NPTX2, 30% vs. 29% after removing tau PET; **Extended Data Fig. 4e**).

We next examined CSF YWHAG:NPTX2 (SomaScan) levels in dementia with Lewy bodies (DLB; n=29), Parkinson’s disease (PD; n=28), and CU (n=28) from the PDBP cohort (**Extended Data Fig. 5a, Supplementary Table 1**) to inform the generalizability of CSF YWHAG:NPTX2 to additional neurodegenerative syndromes with distinct pathophysiology from FTLD and AD. Compared to CU, PD cases on average exhibited modest elevations in CSF YWHAG:NPTX2 (age- and sex-adjusted β = 0.58 [0.15–1.01] s.d., p = 8.9x10^−3^), whereas DLB cases exhibited marked elevations (age- and sex-adjusted β = 1.52 [1.03–2.01] s.d., p = 2.3x10^−8^) similar to FTLD and AD (**Supplementary Table 2, Extended Data Fig. 5b**). CSF YWHAG:NPTX2 was tightly coupled to Montreal Cognitive Assessment (MoCA) total scores across the total cohort (*r* = -0.73, *p* = 3.09x10^−15^) and within PD cases (*r* = -0.41, *p* = 3.19x10^−2^), suggesting stronger YWHAG:NPTX2 elevations in patients with more widespread cortical involvement (**Extended Data Fig. 5c-e)**.

### CSF YWHAG:NPTX2 increases with presymptomatic and symptomatic familial FTLD

We next examined data from the ALLFTD and GENFI cohorts (**Supplementary Table 3**) to test whether CSF YWHAG:NPTX2 was altered in familial FTLD. Both cohorts included healthy non-carrier controls and *C9orf72*, *GRN*, and *MAPT* mutation carriers, encompassing both presymptomatic and symptomatic stages (**Extended Data Fig. 6a-b)**. ALLFTD CSF YWHAG:NPTX2 data were generated using the SomaScan assay (n=249), which exhibited high cross-platform concordance in a subset of 46 samples with paired multiple reaction monitoring mass spectrometry (MRM-MS; *r*=0.92, **Extended Data Fig. 6c**). GENFI data were generated by tandem mass tag mass spectrometry (TMT-MS; n=214).

Because SomaScan (ALLFTD) and TMT-MS (GENFI) report CSF YWHAG:NPTX2 on different arbitrary scales, their values are not directly comparable. To partially overcome this limitation, we normalized values within each cohort as standard deviations from its own healthy controls, which exhibited similar age distributions across cohorts (**Supplementary** Fig. 1a). After normalization, values from control and presymptomatic carrier distributions overlapped closely between cohorts (**Supplementary** Fig. 1b-c). We then residualized values against the healthy control-based age regression line, since symptomatic carriers were substantially older than presymptomatic carriers and non-carriers (**Extended Data** Fig. 6b, **Supplementary** Fig. 1d). This healthy reference z-scored, age-adjusted metric was used for all downstream cross-sectional analyses.

We found age-adjusted CSF YWHAG:NPTX2 levels were markedly elevated in symptomatic familial FTLD mutation carriers in each gene group compared to presymptomatic carriers and non-carrier controls healthy subjects, both in ALLFTD and GENFI in regression models controlling for sex (**Fig. 2a-b, Supplementary Table 4**). In ALLFTD, CSF YWHAG:NPTX2 levels were significantly elevated in presymptomatic *MAPT* compared to controls, whereas in GENFI levels were elevated in presymptomatic *MAPT* and *C9orf72* carriers compared to controls.

**Figure 2.**
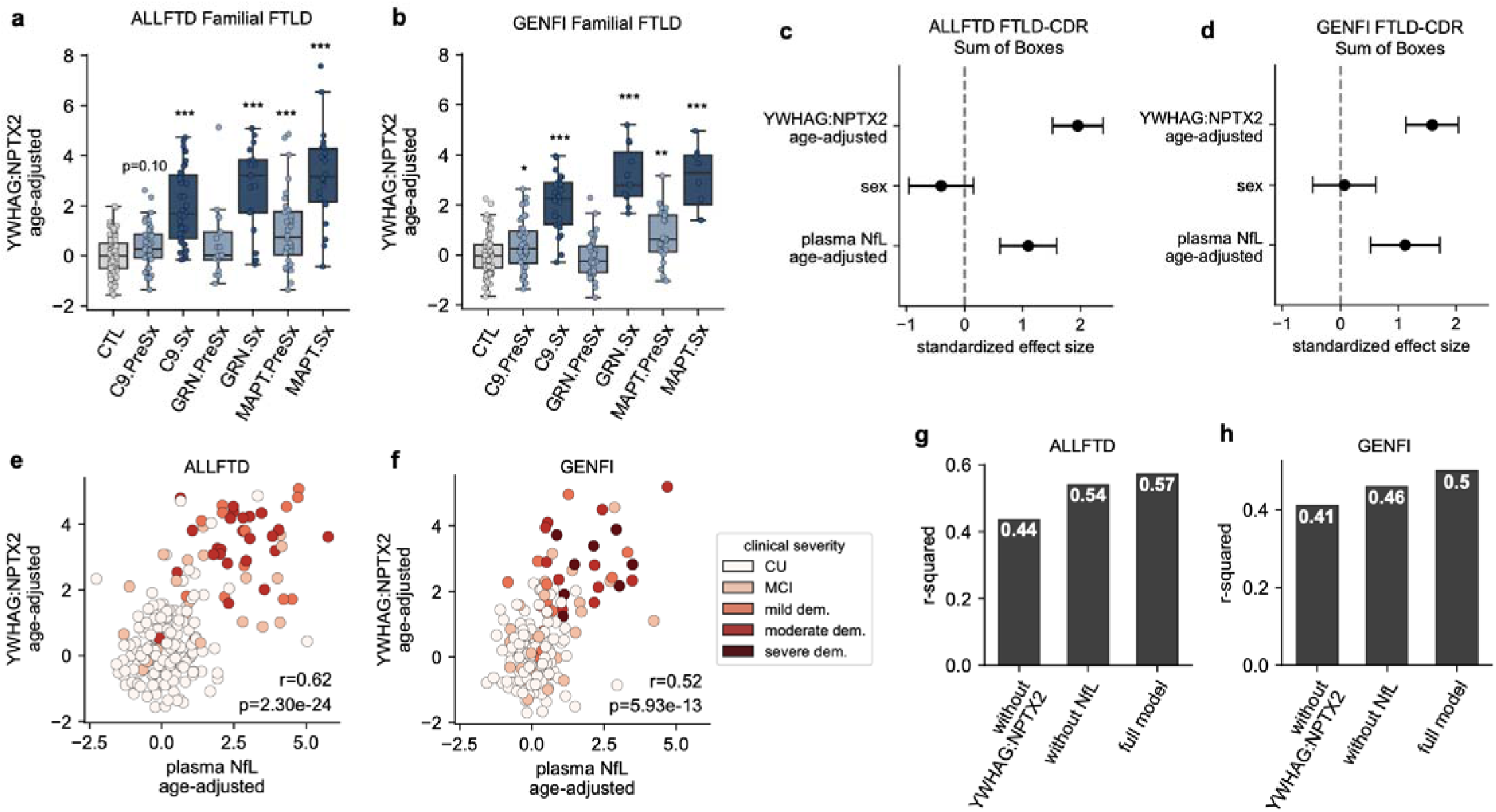
CSF YWHAG:NPTX2 ratio increases with presymptomatic and symptomatic genetic FTLD. **a**, Non-carrier control, presymptomatic (PreSx), and symptomatic (Sx) genetic FTLD stage versus age-adjusted CSF SomaScan YWHAG:NPTX2 in the ALLFTLD cohort (n=249); non-carrier n=83, C9orf72 PreSx n=40 / Sx n=38, GRN PreSx n=17 / Sx n=17, MAPT PreSx n=32 / Sx n=22). The box bounds are the Q1, median and Q3; the whiskers show Q1 − 1.5× the interquartile range (IQR) and Q3 + 1.5× the IQR. **b,** As in **a**, but for the GENFI cohort (n=214) with CSF mass-spec YWHAG:NPTX2; non-carrier n=64, C9orf72 PreSx n=44 / Sx n=24, GRN PreSx n=38 / Sx n=11, MAPT PreSx n=25 / Sx n=8). **c,** CDR+NACC-FTLD (FTLD-CDR) Sum of Boxes regressed against age-adjusted CSF YWHAG:NPTX2, sex, and age-adjusted plasma NfL (Quanterix Simoa) in a linear model in the ALLFTLD cohort (n=217). Points and error bars represent standardized effect sizes and 95% confidence intervals. **d,** As in **c,** but in the GENFI cohort (n=165). **e,** Linear correlation and p-value between age-adjusted CSF YWHAG:NPTX2 and age-adjusted plasma NfL (log_10_) in the ALLFTLD cohort, color-coded according to FTLD-CDR Global (n=217). **f,** As in **e**, but in the GENFI cohort (n=165). **g,** R-squared results from a linear model regressing FTLD-CDR Sum of Boxes against covariates displayed on the x-axis in the ALLFTLD cohort (n=217). Full model includes age-adjusted CSF YWHAG:NPTX2, sex, and age-adjusted plasma NfL. **h,** as in **g**, but in the GENFI cohort (n=165).

Similar to sporadic FTLD, CSF YWHAG:NPTX2 showed a robust association with clinical severity in ALLFTD (n=217, standardized β = 1.95 [1.52–2.39], p = 2.9x10^−16^) and GENFI (n=165, standardized β = 1.59 [1.14–2.04], p = 8.9x10^−11^), independent of plasma NfL (Quanterix Simoa), and sex (**Fig. 2c-d**). Plasma NfL showed a smaller but significant independent association (ALLFTD standardized β = 1.10 [0.61–1.59], p = 1.3x10^−5^; GENFI standardized β = 1.12 [0.52–1.72], p = 3.0x10^−4^) compared to CSF YWHAG:NPTX2, again similar to sporadic FTLD (**Fig. 2c-d**). Visualization of CSF YWHAG:NPTX2 versus plasma NfL showed moderate correlation in both familial FTLD cohorts (ALLFTD r=0.62; GENFI r=0.52; **Fig. 2e-f**). Comparing the clinical severity percent variance explained by full and reduced models, removing CSF YWHAG:NPTX2 reduced variance explained by 14% (57% à 44%) in ALLFTD and 9% (50% à 41%) in GENFI, while removing plasma NfL reduced it by 3% (57% à 54%) in ALLFTD and 4% (50% à 46%) in GENFI (**Fig. 2g-h**).

We also conducted sensitivity analyses in a subset of 109 ALLFTD participants with available immunoassay-based CSF NfL (Quanterix Simoa). CSF YWHAG:NPTX2 was moderately correlated with CSF NfL (*r*=0.47, **Extended Data Fig. 6d)** and exhibited a stronger association with clinical severity than CSF NfL (**Extended Data Fig. 6e-f)**, consistent with the larger plasma NfL models. Collectively, these data show CSF YWHAG:NPTX2 is a robust independent correlate of cognitive impairment in familial FTLD, similar to sporadic FTLD.

### CSF YWHAG:NPTX2 is estimated to rise prior to NfL and symptom onset in familial FTLD

We observed significant but subtle group-level differences in CSF YWHAG:NPTX2 between some presymptomatic familial FTLD gene groups and controls. Since the presymptomatic period spans multiple decades, we next asked whether CSF YWHAG:NPTX2 levels varied as a function of proximity to symptom onset. We leveraged individualized disease progression models, previously developed using longitudinal clinical, neuropsychological, regional brain MRI, and plasma biomarker data in ALLFTD and GENFI, to estimate years before or after symptom onset^2^. Plotting CSF YWHAG:NPTX2 against estimated years from symptom onset, or “disease age”, we observed that CSF YWHAG:NPTX2 began to rise in presymptomatic carriers compared to controls prior to estimated symptom onset in both cohorts (**Fig. 3a**), with consistent patterns across mutation types (**Extended Data Fig. 7a-b**).

**Figure 3.**
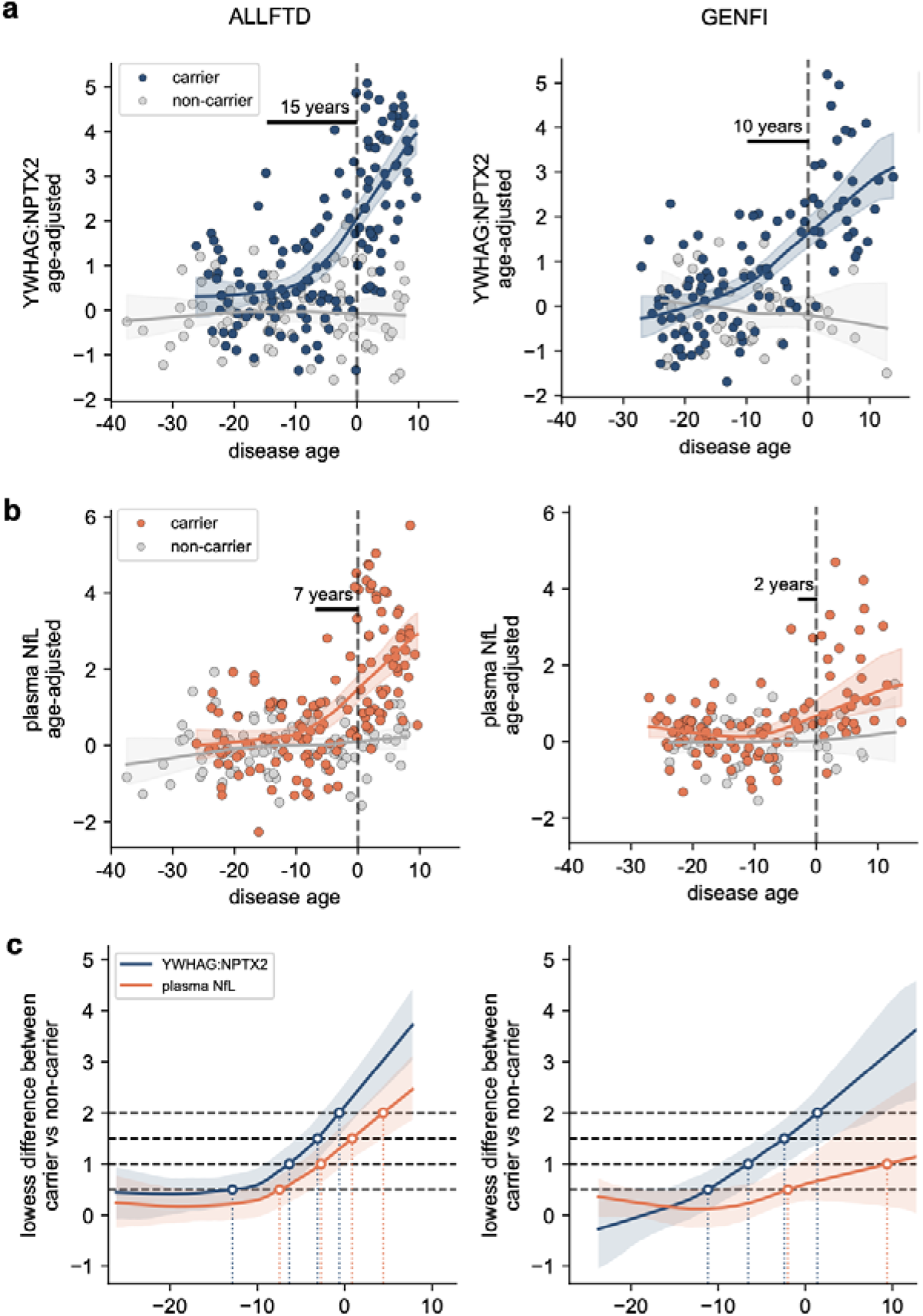
CSF YWHAG:NPTX2 is estimated to rise prior to NfL and symptom onset in familial FTLD. **a**, Changes with estimated years until symptom onset of age-adjusted CSF YWHAG:NPTX2 in the ALLFTLD (n=217; 77 non-carriers, 140 carriers; left) and GENFI (n=150; 44 non-carriers, 106 carriers; right) cohorts. Lowess regression lines with 95% confidence interval for mutation carriers (orange) versus non-carriers (blue) are shown. **b,** As in **a**, but for age-adjusted plasma NfL. **c,** Carrier-minus-non-carrier difference in each marker across disease age in ALLFTD (left) and GENFI (right), with the ages at which the difference first reaches 0.5, 1.0, 1.5, 2.0 standard deviations marked. Bands are 95% bootstrapped confidence intervals.

To estimate how early CSF YWHAG:NPTX2 began to increase, we determined the point at which the lowess regression lower 95% confidence interval of carriers diverged from the upper 95% confidence interval of controls, in line with prior approaches (**Supplementary Table 5**)^20,21^. These analyses estimated CSF YWHAG:NPTX2 to rise 15 years before symptom onset in ALLFTD (n=217) and 10 years in GENFI (n=150). Conversely, plasma NfL was estimated to increase 7 years before estimated symptom onset in ALLFTD and 2 years in GENFI (Fig. 3b). CSF NfL was similarly estimated to increase 2 years before symptom onset in ALLFTD (**Extended Data Fig. 6g**), though this analysis was limited by a smaller available sample (n=97).

Because these divergence points depend on how precisely the confidence interval for each curve is estimated, which is sensitive to sample size, we next investigated divergence based on effect size benchmarks. For each marker, we determined the disease age at which the difference between carriers and non-carriers first reached 0.5, 1.0, 1.5 and 2.0 standard deviations (**Supplementary Table 5**). By this metric, CSF YWHAG:NPTX2 reached every threshold several years earlier than plasma NfL. CSF YWHAG:NPTX2 reached 0.5 s.d approximately 12.9 years before estimated onset in ALLFTD and 11.2 years in GENFI, compared with 7.5 and 2.0 years for plasma NfL (**Fig. 3c**). At 1.0 s.d., CSF YWHAG:NPTX2 reached the threshold 6.3 and 6.6 years before onset, respectively, whereas plasma NfL reached it 2.7 years before onset in ALLFTD and only after symptom onset in GENFI (Fig. 3c, Supplementary Table 5). Because plasma NfL contributed to the model used to estimate disease age, we repeated these analyses using disease age estimates fitted without plasma NfL as an input in ALLFTD (n=188). Divergence estimates based on confidence interval and effect-size benchmarks were essentially unchanged (**Extended Data** Fig. 7c).

Though precise estimations differ slightly between cohorts, likely due to proteomics platform and patient composition, overall our data show that CSF YWHAG:NPTX2 elevations capture preclinical disease progression in FTLD prior to rises in NfL, possibly a decade or more before symptom onset.

### CSF YWHAG:NPTX2 and plasma NfL together predict future clinical conversion in presymptomatic familial FTLD

Biomarkers that predict future phenoconversion among presymptomatic individuals can support clinical trial enrichment by identifying individuals most likely to decline, and therefore benefit from treatment, during the trial period^5,9,10,22^. Building on our disease progression models, we next asked whether CSF YWHAG:NPTX2, alone or in combination with NfL, could prospectively predict clinical conversion. We analyzed non-age-adjusted values to mirror clinical context. We analyzed longitudinal clinical data with up to 8 years of follow-up in ALLFTD and up to 5 years of follow-up in GENFI (**Extended Data** Fig. 8a-b). In ALLFTD participants with CSF YWHAG:NPTX2 and available plasma NfL, 21 of 75 presymptomatic mutation carriers at CSF draw (CDR®+NACC-FTLD Global = 0) phenoconverted during the follow-up period (CDR®+NACC-FTLD Global ≥ 0.5). Phenoconverters and non-converters had an equivalent mean follow-up duration of 5.5 years. In GENFI, 13 of 46 presymptomatic mutation carriers at CSF draw phenoconverted during the follow-up period, and phenoconverters had on average 1-year longer follow-up compared to non-converters (3.3 years versus 2.3). Progression to more severe impairment was less frequent: 5 of the 75 ALLFTD carriers reached a CDR®+NACC-FTLD-SB above 10 during follow-up, and only 1 of 46 in GENFI (**Extended Data** Fig. 8a-b).

We next tested whether binary biomarker cut-offs could accurately predict phenoconversion. Because a prediction threshold depends on the horizon over which phenoconversion is expected to occur, we computed IPCW-weighted time-dependent ROC curves across the follow-up period in ALLFTD (**Extended Data Fig. 8c**). No events were detected at 1 year follow-up. Both biomarkers were most predictive of phenoconversion at 1.5 years (CSF YWHAG:NPTX2 AUC = 0.93; plasma NfL AUC = 0.84), a duration that falls within the window of a typical dementia prevention clinical trial^23,24^. At 1.5 years follow-up, we selected the log10-plasma NfL threshold at ≥80% sensitivity (minimizing false negatives, appropriate for a screening biomarker) and the CSF YWHAG:NPTX2 threshold at ≥90% specificity (minimizing false positives, appropriate for a confirmatory CSF biomarker), classifying individuals into four biomarker groups (**Extended Data Fig. 8d**). Deriving thresholds at longer horizons made both markers more inclusive, but the values remained within a narrow range (**Extended Data Fig. 8e**). Interestingly, the resulting SomaScan CSF YWHAG:NPTX2 threshold (0.24) was only 0.19 z-scores higher than the threshold we previously established (0.05) for distinguishing MCI from healthy controls and predicting future phenoconversion in amyloid-positive healthy controls^16^ (**Extended Data Fig. 8f**). These thresholds are directly comparable because CSF YWHAG:NPTX2 was measured on the same proteomic platform (v.4.1 SomaScan), using the same aptamers and standardized against the same fixed reference means and s.d. constants as in that study rather than re-standardized within the present cohort.

Applying these groups, we plotted CSF YWHAG:NPTX2 versus plasma NfL among ALLFTD presymptomatic mutation carriers at baseline, with points color-coded by eventual conversion to symptomatic disease within 1.5 years and size-coded according to time until symptom onset or last follow-up if censored. This revealed that individuals positive for both plasma NfL and CSF YWHAG:NPTX2 were highly enriched for imminent phenoconversion (**Fig. 4a**). We next examined Kaplan-Meier curves across the four biomarker groups. The double-positive (+/+) group declined rapidly with 6 of 11 carriers phenoconverting within 1.5 years, whereas no carriers positive for plasma NfL alone (+/–) converted in that window (**Fig. 4b**). Over the full 8-year follow-up, Cox proportional hazard regression covarying for baseline age, sex, and education confirmed a 17-times increased risk of symptom conversion in the +/+ group (hazard ratio, HR = 17.18 [3.74–78.91], p = 2.6x10–4) relative to the –/– reference group, while +/– carriers showed no clear increase in risk (HR = 1.83 [0.24–13.94], p = 0.56) (Fig. 4c, Supplementary Table 6). Parallel analyses using CSF YWHAG or CSF NPTX2 individually confirmed that the ratio provided the best discrimination of future converters (**Extended Data Fig. 9a-c**). Substituting CSF NfL for plasma NfL produced a similar pattern of results (**Extended Data Fig. 9d-e, Supplementary Table 7**).

**Figure 4.**
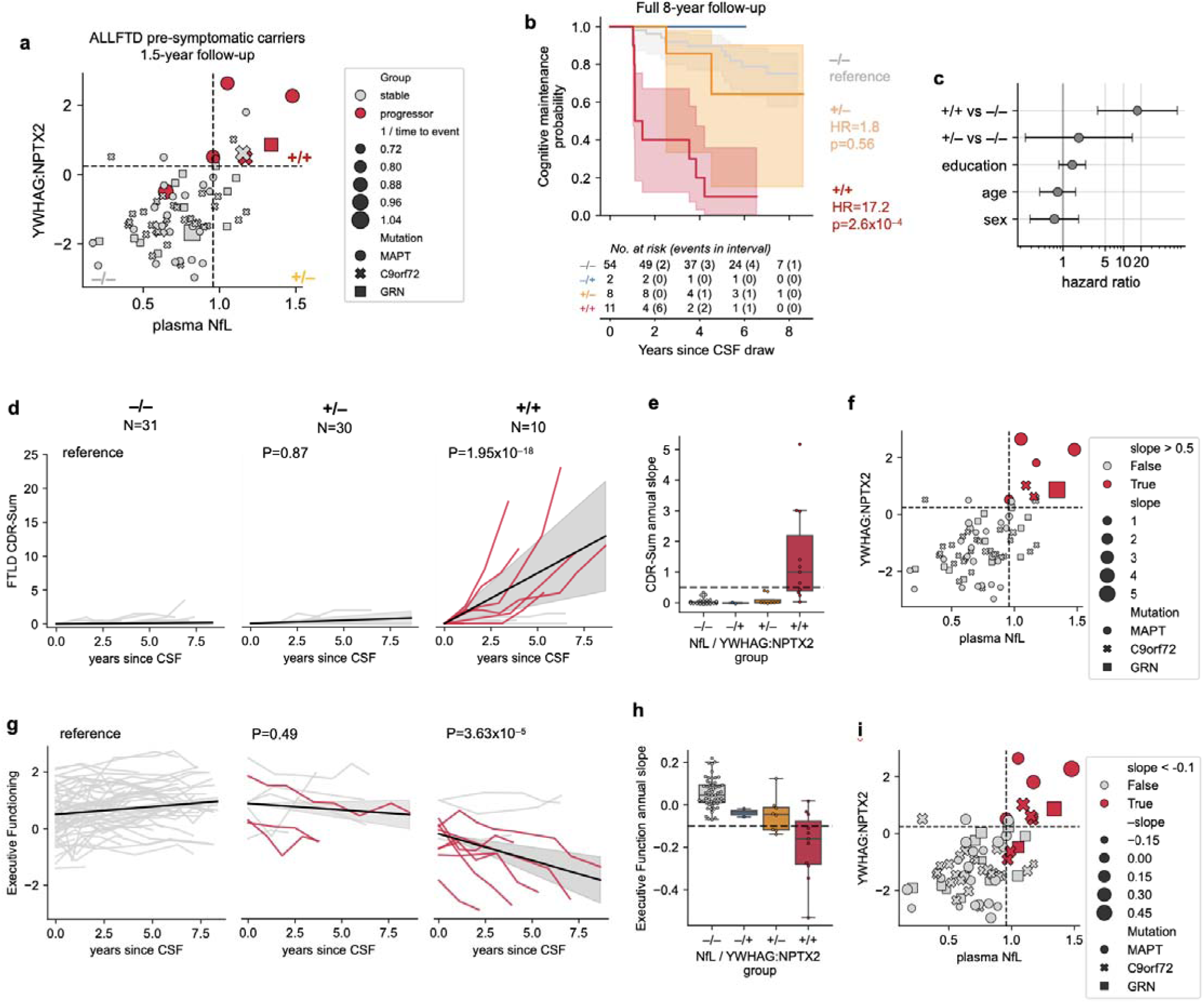
CSF YWHAG:NPTX2 ratio and plasma NfL together predict actual symptom onset in presymptomatic genetic FTLD. **a**, Scatterplot showing CSF YWHAG:NPTX2 versus plasma NfL in presymptomatic genetic FTLD in the ALLFTLD cohort, with points color-coded by symptomatic conversion within 1.5 follow-up years, sized by time until conversion, and symbolized by mutation status (n=75; –/– n=54, –/+ n=2, +/– n=8, +/+ n=11). Dotted lines define positive and negative biomarker gates determined by 80% sensitivity for plasma NfL and 90% specificity for CSF YWHAG:NPTX2. **b,** Kaplan Meier curves with 95% confidence intervals showing symptom onset over 8 follow-up years in presymptomatic genetic FTLD, stratified by biomarker groups defined by gates in **a**. Hazard ratios and p-values from the Cox proportional hazard model in **c** and the sample sizes per group and number of events per year are shown. **c,** Results from a Cox proportional hazard model regressing time until symptom conversion against biomarker groups relative to the double negative (–/– reference group), education, age, and sex. Points and error bars represent effect sizes and 95% confidence intervals. **d,** Spaghetti plot showing longitudinal trajectories in CDR+NACC-FTLD (FTLD CDR) Sum of Boxes for all individuals, stratified by biomarker group over 8 follow-up years. Average slopes with 95% confidence intervals are shown. A linear mixed-effects model regressing FTLD CDR-Sum of Boxes against time, biomarker group, and their interaction, covarying for baseline age, sex, education, and with random intercept and slope was tested. P-values from the biomarker group x time interaction term are shown. **e,** Box plot showing biomarker group versus longitudinal slope of FLTD CDR-Sum of Boxes. The box bounds are the Q1, median and Q3; the whiskers show Q1 − 1.5× the interquartile range (IQR) and Q3 + 1.5× the IQR. **f**, Scatterplot showing CSF YWHAG:NPTX2 versus plasma NfL in presymptomatic genetic FTLD in the ALLFTLD cohort, with points color-coded and sized by longitudinal slope of FLTD CDR-Sum of Boxes, and symbolized by mutation status. Dotted lines define positive and negative biomarker gates determined in **a**. **g-i,** As in **d-f,** but for longitudinal executive functioning (Uniform Data Set v3.0) composite scores.

We next tested a linear mixed-effects model to assess whether YWHAG:NPTX2 and NfL biomarker-defined groups in ALLFTD exhibited different slopes of clinical progression over time (**Supplementary Table 8**). We regressed CDR®+NACC-FTLD-SB against time, biomarker group, and their interaction, covarying for baseline age, sex, education, with random intercepts and slopes per subject. Notably, the +/+ group showed a substantially steeper increase in clinical severity over time (β = 1.47 [1.04–1.89], p = 1.2x10^−11^), with an average annual increase of 1.5 points relative to the –/– group (**Fig. 4d**). Pronounced slopes of clinical progression (annual CDR®+NACC-FTLD-SB slope > 0.5) were completely restricted to the +/+ group (**Fig. 4d-f**). 7 of 19 plasma NfL+ individuals (37%) and 7 of 11 +/+ individuals (64%) showed a CDR®+NACC-FTLD-SB annual slope greater than 0.5. In other words, using both plasma NfL and CSF YWHAG:NPTX2 identified fast clinical progressors 1.7-fold compared to plasma NfL alone. We next analyzed longitudinal mixed effect models using the NACC Uniform Data Set (V.3.0) executive function composite score (UDS3-EF), a continuous outcome sensitive to early cognitive deficits in familial FTLD^25,26^. For UDS3-EF, we observed a similar pattern by which the +/+ group exhibited significantly steeper declines in cognition (β = –0.23 [–0.32 to –0.15], p = 4.7x10^−8^) relative to –/– individuals, with the +/– group showing an intermediate significant decline (β = –0.10 [–0.19 to –0.01], p = 0.04) (**Fig. 4g-i**, **Supplementary Table 9**). Model residual diagnostics confirmed normality assumptions were met for UDS3-EF, while minor non-normality in CDR®+NACC-FTLD-SB residuals was expected given its bounded (**Supplementary Fig. 2a-b**), ordinal distribution and is consistent with standard practice in AD clinical trials^23,24^.

We next tested the ALLFTD-derived thresholds in the GENFI cohort. Because the two cohorts measured CSF YWHAG:NPTX2 on different platforms and plasma NfL with different Quanterix kit versions, we re-expressed both thresholds (**Extended Data Fig. 10a**) on the healthy-control z scale described previously (**Supplementary Fig. 1**), here without age adjustment in order to mirror clinical practice. Applying the ALLFTD-derived thresholds to GENFI proved stringent with only a single double-positive (+/+) carrier, with the remainder falling into the –/– (n=41) or +/– (n=4) groups (**Extended Data Fig. 10b**). Despite the small number, this +/+ individual phenoconverted within 1.2 years of CSF draw and exhibited by far the most rapid and severe clinical decline among all GENFI presymptomatic carriers during follow-up (final CDR®+NACC-FTLD-SB = 12.5; **Extended Data Fig. 8b**, **Extended Data Fig. 10b-e**). While the single double-positive case limits formal statistical inference, the magnitude and speed of this individual’s deterioration is consistent with the ALLFTD finding that double positivity enriches for imminent phenoconversion and steep decline.

Finally, we asked how adding CSF YWHAG:NPTX2 to a plasma NfL screen would affect patient selection for a prevention trial. Of 75 presymptomatic carriers in ALLFTD, 19 (25%) were plasma NfL-positive and would be selected by that screen alone; 11 of these were also CSF YWHAG:NPTX2-positive (**Supplementary Fig. 3a**). The +/+ individuals converted markedly faster (**Supplementary Fig. 3b**), and all 6 phenoconversions within 1.5 years occurred among them. This results in an event rate of 55% when considering both biomarkers compared to 32% for plasma NfL positivity alone (1.7-fold increase, **Supplementary Fig. 3c**). Under standard clinical trial assumptions of a 50% relative reduction in phenoconversion, 80% power and two-sided α = 0.05, a trial enrolling on plasma NfL alone would require 225 participants to observe a significant effect, whereas a trial enrolling on both biomarkers would require only 100, a 2.3-fold reduction (bootstrap 95% CI 1.4–4.8) that was stable across the plausible range of treatment effects (**Supplementary Fig. 3d-e**). Because only plasma NfL-positive carriers would undergo lumbar puncture, the two-tiered strategy would also require fewer blood tests (679 versus 889) at the cost of 172 CSF draws (**Supplementary Fig. 3f**). Though these projections are likely optimistic given the small sample size and lack of a well-powered validation cohort, they illustrate potential benefit of measuring CSF YWHAG:NPTX2 for FTLD prevention trials.

Together, these data show that CSF YWHAG:NPTX2, especially when combined with the minimally invasive predictive biomarker plasma NfL, dramatically improves prediction of imminent clinical conversion and subsequent cognitive progression in presymptomatic familial FTLD, with potential to improve clinical trial design.

## Discussion

Our study establishes the CSF YWHAG:NPTX2 ratio as a cross-dementia biomarker candidate with robust associations with clinical severity across multiple sporadic clinical subtypes and familial forms of FTLD. CSF YWHAG:NPTX2 relationships with clinical severity were stronger and independent of NfL, the most commonly used fluid biomarker for FTLD assessment. We also extend prior observations in AD to atypical AD variants with similar ages-of-onset to FTLD, underscoring the potential of CSF YWHAG:NPTX2 to capture shared synaptic underpinnings of cognitive decline across early-onset dementias. Strikingly, CSF YWHAG:NPTX2 increases ∼10-15 years before estimated symptom onset in presymptomatic FTLD gene carriers and together with plasma NfL strongly predicts imminent phenoconversion, positioning it as a promising prognostic tool for clinical trial participant stratification^10^. This suggests that a two-tiered screening strategy beginning with plasma NfL followed by CSF YWHAG:NPTX2 (akin to plasma pTau217 and tau PET in AD^27^) could markedly improve patient selection for prevention trials in familial FTLD. Moreover, CSF YWHAG:NPTX2 may be a useful clinical endpoint to assess treatment-related changes in synaptic integrity linked to cognitive function^28^. Validation of these strategies should be explored in future studies.

The strong association between CSF YWHAG:NPTX2 and disease severity was consistent across sporadic and autosomal dominant FTLD groups, underscoring its potential as a robust marker of clinical progression regardless of underlying etiology. This is particularly relevant in sporadic FTLD, where validated proteinopathy-specific biomarkers are not currently available to guide diagnosis and progressive proteinopathy burden. As novel molecular biomarkers targeting specific proteinopathies (e.g., FTLD-TDP or FTLD-tau subtypes) continue to develop, it will be important to evaluate whether YWHAG:NPTX2 retains its value as a disease-agnostic synaptic marker of clinical severity. Importantly, our findings across FTLD, AD, and DLB suggest that YWHAG:NPTX2 is more tightly coupled to cognitive status than any given proteinopathy-specific measure or syndromic group, supporting its continued relevance as a nonspecific biomarker of cognitive risk and resilience.

Interestingly, the early rise of CSF YWHAG:NPTX2 in presymptomatic familial FTLD mirrors its trajectory in autosomal dominant AD, where it increases 10-20 years before symptom onset^16^. This consistency across AD and FTLD underscores synaptic dysfunction as a common early feature of these dementias and a potential therapeutic target for dementia prevention. NPTX2 is a synapse-specific protein that is directly perturbed by tau and TDP-43 pathology^29,30^, and its reduction in presymptomatic disease may reflect early loss of inhibitory–excitatory synaptic integrity. Further, overexpression of NPTX2 in brains of TauP301S mice prevents tau-induced synapse loss, suggesting beneficial roles of NPTX2 in preserving synaptic function^30^. YWHAG (14-3-3γ) is a broadly expressed neuronal regulatory protein that coordinates cellular stress responses, trafficking, and survival pathways^31,32^. YWHAG is also implicated in proteinopathies, including regulation of TDP-43 nuclear–cytoplasmic shuttling^33^ and tau-related stress signaling^34^. In combination, YWHAG and NPTX2 may capture both early-stage neuronal stress and synaptic failure that are not fully captured by axonal loss markers such as NfL. Future studies should clarify the biological basis of CSF YWHAG and NPTX2 alterations, including their relationships to synaptic loss, neuronal stress responses, and the accumulation of amyloid, tau, TDP-43, and synuclein pathology in AD and FTLD.

We previously demonstrated that CSF YWHAG:NPTX2 explained more variance in cognitive impairment than Aβ, tau, and NfL in predominantly late-onset AD and rose prior to estimated symptom onset in autosomal dominant AD^16^. Here, we extend these observations into sporadic, early-onset variants of AD, which are typically characterized by greater tau PET burden than late-onset AD, irrespective of clinical phenotype^35^. In our early-onset enriched UCSF-MAC cohort, we found that CSF YWHAG:NPTX2 correlated more strongly with clinical severity than both fluid-based and PET-based measures of amyloid and tau pathology. Notably, this was observed in a rigorously phenotyped sample with tau PET and CSF collected within six months. We similarly observed elevated YWHAG:NPTX2 in DLB and PD cases with cognitive impairment. These findings suggest that YWHAG:NPTX2, as a synaptic measure, more directly reflects the clinical sequelae of pathogenic protein accumulation such as tau and alpha-synuclein. Future work should examine how longitudinal changes in YWHAG:NPTX2 relate to concurrent changes in fluid biomarkers of AD tau pathophysiology including MTBR243 tau^11^, which more directly reflects insoluble tau aggregate deposition, and phosphorylated tau species like pTau217 and pTau231, which capture earlier effects of beta amyloid plaque accumulation on tau.

Our study has several limitations. Although we included longitudinal clinical data and observed robust associations with clinical severity across multiple contexts, the sample sizes in some subgroups such as presymptomatic FTLD phenoconverters, specific mutation groups (i.e. *MAPT, GRN*), and MND, remain modest, necessitating replication in larger samples with longer follow-up. In particular, only three presymptomatic GRN carriers phenoconverted during follow-up across ALLFTD and GENFI, which likely limited power to detect CSF YWHAG:NPTX2 alterations within presymptomatic GRN. Additional samples and studies may also reveal the value of differential weighting of the two proteins for different use cases (disease staging vs early prognosis). Future studies should also analyze CSF YWHAG:NPTX2 from longitudinal CSF collections, autopsy validated FTLD samples, vascular dementia, non-degenerative neurological injury (e.g., stroke, epilepsy), and more racially/ethnically diverse cohorts will help address specificity to dementia and neurodegeneration and further enhance generalizability. While our use of multiple cohorts and proteomic techniques, including aptamer-based and mass spectrometry approaches, enhances generalizability and mitigates platform-specific biases, the current reliance on large-scale proteomics limits immediate clinical translation. Development of a targeted, clinically deployable assay for CSF YWHAG:NPTX2 with high reproducibility and validated cut-offs will be essential for synaptic ratio biomarker development in real-world settings. This will likely require development of reliable CSF YWHAG and NPTX2 enzyme-Linked Immunosorbent Assay (ELISA) assays that can be adopted onto more advanced ELISA-based platforms (i.e. Lumipulse, Simoa, Elecsys). While high-performing assays for NPTX2 are available^36,37^, assays for YWHAG are absent, to our knowledge. Future studies should also evaluate whether YWHAG:NPTX2 can be reliably measured in plasma, which could enable more scalable and minimally invasive applications.

In conclusion, the CSF YWHAG:NPTX2 ratio is a powerful predictor of clinical progression in FTLD, and its use in combination with plasma NfL could support more precise disease staging and prognosis in clinical trials across the FTLD clinico-pathological continuum.

## METHODS

### Cohorts

Cohort demographic sample sizes are provided in **Supplementary Tables 1 and 3**. All cohorts were sex balanced.

### UCSF Memory and Aging Center (MAC)

Participants included 144 patients with sporadic forms of FTLD, 149 patients with sporadic AD, and 86 CU controls enrolled in research studies at the UCSF Memory and Aging Center (MAC). All participants underwent lumbar puncture for collection of CSF samples, genetic screening, and comprehensive clinico-diagnostic procedures, including neurological examination, cognitive testing, and informant interview. The UCSF Institutional Review Board (IRB) approved study procedures, and all participants provided written informed consent or assent with proxy consent.

Clinical diagnoses were made via multidisciplinary consensus case conference with board-certified neurologists and neuropsychologists. FTLD cases met published criteria for a primary FTLD clinical syndrome, specifically behavioral variant FTD (bvFTD)^38^ with or without motor neuron disease (MND)^39^, semantic variant primary progressive aphasia (svPPA)^40^, nonfluent variant PPA (nfvPPA)^40^, corticobasal syndrome with negative AD biomarkers (CBS-FTD)^41^, or progressive supranuclear palsy spectrum syndromes (PSPs)^42^. All AD cases met criteria for cognitive impairment^43,44^ and had at least one positive AD biomarker based on available CSF pTau181:Aβ42 ratios or amyloid PET visual read. UCSF-MAC AD cases were enriched for early-onset clinical variants with a comparable age distribution to FTLD cases. Specifically, AD cases met published criteria for early-onset amnestic AD (onset before age 65 years; EOAD; n=47)^45^, posterior cortical atrophy (PCA; n=23)^46^, logopenic variant primary progressive aphasia (lvPPA; n=24)^40^, behavioral variant AD (bvAD; n=6)^47^, CBS due to AD (CBS-AD; n=13)^41^, mild cognitive impairment due to AD (MCI-AD; n=22)^44^, or late-onset amnestic AD (onset after age 65 years; LOAD; n=13)^43^. CU controls were functionally intact (CDR global=0), community-dwelling volunteers classified as clinically normal at consensus review. All participants were free of major neuromedical confounds (e.g., HIV, brain tumor) and screened negative for autosomal dominant FTLD or AD mutations.

### ARTFL/LEFFTDS Longitudinal Frontotemporal Lobar Degeneration (ALLFTD) Consortium

Participants included 166 carriers of pathogenic mutations in the *C9orf72* (n=78)*, GRN* (n=34) or *MAPT (n=54)* genes and 83 noncarrier controls from families with a known mutation in one of these genes^48^. Participants were enrolled in the ALLFTD consortium (<u>NCT04363684</u>)^9,49^, which includes 27 collaborating centers across the US and Canada. The ALLFTD study was approved through the Trial Innovation Network at Johns Hopkins University. Local ethics committees at each of the sites approved the study, and all participants provided written informed consent or assent with proxy consent. Inclusion in the present study required completion of baseline lumbar puncture for CSF collection

### Genetic Frontotemporal Dementia Initiative (GENFI)

To validate familial FTLD models across cohorts and proteomic platforms, we leveraged publicly-available CSF tandem mass tag (TMT) mass spectrometry YWHAG:NPTX2 data from the GENFI cohort^18^. The GENFI CSF TMT mass spec dataset included 68 *C9orf72,* 49 *GRN*, and 33 *MAPT* carriers as well as 64 noncarrier controls, drawn from 14 centers distributed across Europe and Canada. The London Queen Square Ethics committee and local ethics committees at each site approved GENFI study protocols, which complied with the Declaration of Helsinki. All participants provided written informed consent at enrollment.

### Parkinson’s Disease Biomarker Program (PDBP)

CSF samples were acquired from participants enrolled in the Parkinson’s Disease Biomarker Program (PDBP), a multi-site, NIH-funded biorepository focused on PD-spectrum disorders. DLB and PD cases were diagnosed at each site according to established clinical criteria^50,51^. A balanced subset of 30 PD, 30 DLB, and 30 CU controls were selected from the available pool of CSF samples with MoCA scores, with the total sample size determined by the proteomic assay budget. No exclusions were made based on biomarker status, Parkinson’s disease medication use, comorbidities, or potential co-pathologies. Five samples were excluded during proteomic quality control based on outlier measurements, resulting in the final analytic sample of 85 participants (29 DLB, 28 PD, and 28 CU controls). Local IRB committees approved PDBP study procedures, including sample collections, and all participants provided written informed consent or assent with proxy consent.

### CSF processing

All studies followed standard collection, processing and storage protocols for CSF samples. CSF was collected in polypropylene tubes via lumbar puncture in lateral recumbent or sitting positions. CSF samples were centrifuged at 2000 g for 10 minutes at room temperature. Supernatant was aliquoted in polypropylene tubes and stored at -80°C until further analyses. All CSF samples underwent no more than one freeze/thaw cycle before analysis.

### Biomarker assays

CSF samples from UCSF-MAC, ALLFTD, and PDBP were assayed on the SomaLogic platform (v.4.1 SomaScan) in a single batch per cohort. CSF SomaScan YWHAG:NPTX2 ratios were calculated based on our published approach of: 1) log_10_-normalization of aptamer levels (YWHAG SeqID=4179-57, NPTX2 SeqID=6521-35), 2) z-score normalization of log_10_ aptamer values using means and s.d. (YWHAG mean = 3.425, s.d. = 0.183; NPTX2 mean = 4.099, s.d. = 0.171), and 3) taking the difference between YWHAG and NPTX2 z-scores. Means and standard deviations were derived from cohorts in our published study to enable direct comparisons^16^. The normalized ratio approach was shown to optimize associations with dementia severity and outperformed either protein alone.

A subset of samples from ALLFTD (n=46) were also assayed via multiple reaction monitoring mass spectrometry (MRM-MS) to quantify a targeted panel of proteins^52^, including proteotypic peptides for NPTX2 (LESLEHQLR, TESTLNALLQR) and YWHAG (DSTLIMQLLR, NVTELNEPLSNEER). MRM-MS YWHAG:NPTX2 ratios in ALLFTD and TMT-MS YWHAG:NPTX2 ratios in GENFI were computed using the same normalization methods, using sample-specific distributions.

In UCSF-MAC, core CSF biomarkers Aβ42, Aβ40, pTau181, and NfL were measured using fully automated Elecsys immunoassays (Roche Diagnostics International Ltd, Rotkreuz, Switzerland), as described previously^53^. CSF pTau181:Aβ42 positivity status was based on a cut-point of >0.022, as previously described and validated against autopsy^53^. In ALLFTD, plasma and CSF NfL were measured using the Quanterix single-molecule array platform (Simoa) NF-Light digital assay on the HD-X instrument, as previously described^9,14^. In GENFI, plasma NfL was measured via Quanterix Simoa Neurology 4-plex A assay.

### Amyloid and tau PET imaging

Amyloid PET imaging in UCSF-MAC was acquired with 11C-Pittsburgh compound B (PIB, n=107), 18F-Florbetapir (n=63), or 18F-Florbetaben (n=22) using PET or PET-CT scanners. Data acquisition and processing were performed using standard procedures^54,55^. Amyloid positivity status was based on visual read, as previously described and validated against autopsy^54,56,57^. Amyloid PET was also quantified in Centiloids^58^ using previously validated methods^59,60^.

Tau PET was acquired with 18F-Flortaucipir on PET-CT scanners (n=84). Data acquisition and processing were performed using standard procedures, using the inferior cerebellar cortex as a reference and extracting Standardized Uptake Value Ratio (SUVR) values based on MR-defined regions using Freesurfer^61^. The average SUVR value in the temporal meta-ROI was calculated as an index of AD tau level, as previously done^61–63^.

### Clinical and cognitive severity outcomes

Clinical severity was determined with the CDR^64^. AD-specific models used the standard CDR sum of boxes score, one of the most common endpoints in AD clinical trials^24^. FTLD-focused models used the CDR plus Behavioral and Language Domains from the National Alzheimer’s Coordinating Center (NACC) FTLD module (CDR®+NACC-FTLD)^65–67^. The CDR®+NACC-FTLD builds on the standard CDR by including domains more affected in FTLD (language and behavior)^65^, and is the most widely used metric for FTLD disease staging and clinical trial outcomes^6,9,23^. In ALLFTD and GENFI, mutation carriers were classified as presymptomatic or symptomatic based on a CDR®+NACC-FTLD Global score of 0 (presymptomatic) or ≥0.5 (symptomatic). CDR®+NACC-FTLD sum of boxes (SB) was examined as a primary outcome in FTLD clinical severity models.

In ALLFTD, the NACC Uniform Data Set (V.3.0) executive function composite score (UDS3-EF) was also examined as an outcome in prognostic models, providing a continuous, cognitive testing-based alternative to the informant report-based CDR®+NACC-FTLD. The UDS3-EF score is sensitive to early executive function deficits in FTLD spectrum diseases, with lower scores representing worse executive functioning^25,26,68^.

### Familial FTLD disease progression models

We previously developed Bayesian disease progression models in the broader ALLFTD and GENFI cohorts to estimate disease age^9^, a latent variable representing the estimated number of years from symptom onset for any given mutation carrier (positive disease age = post-symptom onset; negative disease = pre-symptom onset). The disease progression model jointly modeled 20 clinical, neuropsychological, imaging, and plasma biomarker inputs across 796 mutation carriers and 412 noncarrier family controls from ALLFTD and GENFI. Disease age was inferred for each individual using a Bayesian mixed-effects framework, which included the multimodal clinico-biomarker data and model priors for time from symptom onset. Clinician estimate of disease duration was a prior for symptomatic carriers (based on clinical interviews) and mean age of onset for the corresponding family mutation group was a prior for presymptomatic carriers and non-carrier controls. The model was trained and validated independently in both cohorts and demonstrated consistent progression trajectories, supporting the harmonized application of disease age across ALLFTD and GENFI.

For the present study, we applied the same disease progression modeling framework with a larger number of observations in ALLFTD (N=776) and GENFI (N=935) and an updated set of input variables. Input variables included symptom severity measures (CDR®+NACC FTLD-SB, Revised Self-Monitoring Scale total score, Revised Amyotrophic Lateral Sclerosis Functional Rating Scale total score), neuropsychological test scores (longest digit span forward, longest digit span backward, Trails A, Trails B, semantic fluency, multilingual naming test total score, figure copy, figure recall), volumetric neuroimaging regional composites (frontal, parietal, temporal, medial temporal, occipital, insula, striatum, thalamus, cerebellum), and plasma biomarkers (NfL, GFAP). We leveraged posterior disease age estimates for each subject at the same timepoint as CSF collection to perform pseudo-temporal modeling of CSF YWHAG:NPTX2 in relation to disease age. Lowess regression was used to analyze biomarker trajectories across disease age. The point at which the lowess regression lower 95% confidence interval of mutation carriers exceeded the upper 95% confidence interval of non-carrier controls was used to estimate the time before symptom onset at which biomarkers diverged between FTLD mutation carriers and controls.

### Association analyses

Individuals with missing proteomics/clinical data were excluded, not imputed, to avoid introducing biases that would hinder interpretation of the findings. Multivariable linear regression was used to determine differences in CSF YWHAG:NPTX2 between clinical diagnosis, while controlling for age and sex. Correlation analyses were used to determine associations between CSF YWHAG:NPTX2 and CSF or plasma NfL. Multivariable linear regression was used to determine associations between CSF YWHAG:NPTX2 and clinical severity, independent of covariates, including NfL, age, and sex. Cox proportional hazards regression was used to determine associations between biomarkers and future symptomatic conversion. Linear mixed-effect models were used to determine associations between biomarkers and longitudinal changes in clinical severity. All covariates used in regression analyses are described in the appropriate sections of the main text and figure legends. Standardized betas refer to betas where the predictor variables, but not the outcome variable, are z-scored. Multiple hypothesis testing was not performed in this study, since analyses focused primarily on the CSF YWHAG:NPTX2 ratio and its associations with multiple correlated outcomes (clinical severity, diagnoses, etc). All correlation coefficients reported are Pearson correlations.

### Clinical trial power analysis

Using the 75 ALLFTD presymptomatic carriers with both biomarkers available, we projected enrollment for a hypothetical prevention trial (**Supplementary Fig. 3**). The outcome was phenoconversion within 1.5 years of CSF draw, the horizon of maximum predictive AUC, and event rates with 95% Wilson score intervals were computed for three nested screening strategies: all carriers, plasma NfL-positive, and double-positive. Required sample size was estimated by the two-proportion normal approximation for a two-arm, 1:1 superiority trial at two-sided alpha = 0.05 and 80% power, assuming a 50% relative reduction in phenoconversion, and is reported as the total across both arms. Sample sizes were also computed across assumed reductions of 25-65%. Because the two strategies are nested and share converters, confidence intervals on the fold reduction were obtained by resampling carriers with replacement (5,000 replicates) rather than resampling the two rates independently. Screening burden was the required enrollment divided by the proportion of carriers passing each screen, with lumbar puncture limited to the plasma NfL-positive subset. These projection assume the observed event rates carry over and that follow-up is complete and are illustrative rather than a true trial design.

## Data availability

Raw data are available upon reasonable request with formal applications to each respective study. Proteomic and clinical data from UCSF-MAC can be requested via UCSF at https://memory.ucsf.edu/research-trials/professional/open-science. ALLFTD proteomic and clinical data requests can be completed via ALLFTD at https://www.org/data. Certain data elements may be restricted due to the potential for identifiability in the context of the sensitive nature of the genetic data. PDBP proteomic data is available via UCSF-MAC and clinical data is available through the NINDS Parkinson’s Disease Biomarkers Program Data Management Resources (https://pdbp.ninds.nih.gov/). GENFI proteomic data are available via the Dryad repository (DOI: 10.5061/dryad.r7sqv9snk). GENFI clinical data will be made available by request from any qualified investigator and can be requested through the GENFI website (genfi.org/contact-us-2) or through Dementias Platform UK (portal.dementiasplatform.uk/Apply).

## Code availability

No novel code or algorithms were generated in this study. All analyses were standard practice using python packages: pandas, numpy, scikit-learn, statsmodels, lifelines, matplotlib, seaborn.

## Author Contributions

H.S.-H.O., R.S., and A.L.B. conceptualized the study. H.S.-H.O. led the analyses and produced the figures. H.S.-H.O. and R.S. had full access to the data in the study and take responsibility for the integrity of the data and the accuracy of the data analysis. H.S.-H.O. and R.S. drafted the manuscript. H.S.-H.O., R.S., and A.L.B. supervised the study. J.H.K., B.F.B., H.J.R., J.S.Y. W.W.S., J.C.R., K.B.C., and A.L.B. obtained funding. All authors critically revised the paper for intellectual content. All authors read and approved the final version of the manuscript.

## Acknowledgments

Data collection and dissemination of the data presented in this paper were supported by the ALLFTD Consortium (U19: AG063911 (A.L.B., H.J.R. and B.F.B.), funded by the National Institute on Aging and the National Institute of Neurological Diseases and Stroke) and the former ARTFL and LEFFTDS Consortia (ARTFL: U54 NS092089 (A.L.B., H.J.R. and B.F.B.), funded by the National Institute of Neurological Diseases and Stroke and National Center for Advancing Translational Sciences; LEFFTDS (U01 AG045390 (H.J.R. and B.F.B.)), funded by the National Institute on Aging and the National Institute of Neurological Diseases and Stroke). The paper was reviewed by the ALLFTD Executive Committee for scientific content. Additional funding support to the UCSF-MAC was provided by the National Institutes of Health (NIH) National Institute on Aging (NIA) for Frontotemporal Dementia: Genes, Images, and Emotions: P01 AG019724 (B.L.M.) and UCSF Alzheimer’s Disease Research Center: P30 AG062422 (G.D.R.). We acknowledge the invaluable contributions of the study participants and families as well as the assistance of the support staff at each of the participating sites. This work is also supported by The Alzheimer’s Drug Discovery Foundation (ADDF; A.G, H.S.-H.O.), the Leon Levy Scholarship in Neuroscience (H.S.-H.O.), the Alzheimer’s Association (grant nos. AARF-23-1145318 (R.S.) and AARG-20-683875 (K.B.C.)), New Vision Research (grant no. CCAD 2024-001-1 (R.S.)), the National Institutes of Health Training Grant T32MH135853 (S.J.R.), an American Academy of Neurology, American Brain Foundation, and Association for Frontotemporal Degeneration Fellowship Award (R.S.), the Bluefield Project to Cure FTD, CurePSP, Larry L. Hillblom Foundation (grant nos. 2018-A-006-NET (J.H.K.) and 2018-A-025-FEL (A.M.S.)), the NIH (grant nos. K23AG090757 (R.S.), K23AG084883 (E.W.P.), R01AG038791 (A.L.B.), R01AG032306 (H.J.R.), K23AG059888 (J.C.R.), K23AG073514 (L.V.), P01AG019724 (B.L.M.), P30AG062422 (W.W.S.), R01AG072475 (K.B.C.), K23AG061253 (A.M.S.), R01AG032289 (J.H.K.), R01AG048234 (J.H.K.), K24AG045333 (H.J.R.)) and the Rainwater Charitable Foundation. Samples from the National Centralized Repository for Alzheimer’s Disease and Related Dementias (NCRAD), which receives government support under a cooperative agreement grant (U24 AG21886) awarded by the NIA, were used in this study. The authors acknowledge the invaluable contributions of the study participant and families as well as the assistance of the support staffs at each of the participating sites.

Data and biospecimens used in preparation of this manuscript were obtained in part from the Parkinson’s Disease Biomarkers Program (PDBP) Consortium, supported by the National Institute of Neurological Disorders and Stroke at the National Institutes of Health (U01NS100620; U01NS100610). Investigators include: Roger Albin, Roy Alcalay, Alberto Ascherio, Thomas Beach, Sarah Berman, Bradley Boeve, F. DuBois Bowman, Shu Chen, Alice Chen-Plotkin, William Dauer, Ted Dawson, Paula Desplats, Richard Dewey, Ray Dorsey, Jori Fleisher, Kirk Frey, Douglas Galasko, James Galvin, Dwight German, Steven Gunzler, Lawrence Honig, Xuemei Huang, David Irwin, Un Kang, Kejal Kantarci, Anumantha Kanthasamy, Daniel Kaufer, Horacio Kaufmann, Qingzhong Kong, James Leverenz, Allan Levey, Carol Lippa, Irene Litvan, Oscar Lopez, Jian Ma, Richard Mailman, Lara Mangravite, Karen Marder, Kelly Mills, Nandakumar Narayanan, Laurie Orzelius, Vladislav Petyuk, Judith Potashkin, Liana Rosenthal, Rachel Saunders-Pullman, Clemens Scherzer, Michael Schwarzschild, Tanya Simuni, Andrew Singleton, David Standaert, Debby Tsuang, David Vaillancourt, Jerrold Vitek, David Walt, Andrew West, Cyrus Zabetian, and Jing Zhang. PDBP Investigators not included as co-authors did not participate in reviewing the data analysis or content of the manuscript.

The funding sources had no role in the design and conduct of the study; in the collection, analysis, interpretation of the data; or in the preparation, review or approval of the paper.

---

**Extended Data Figure 1.**
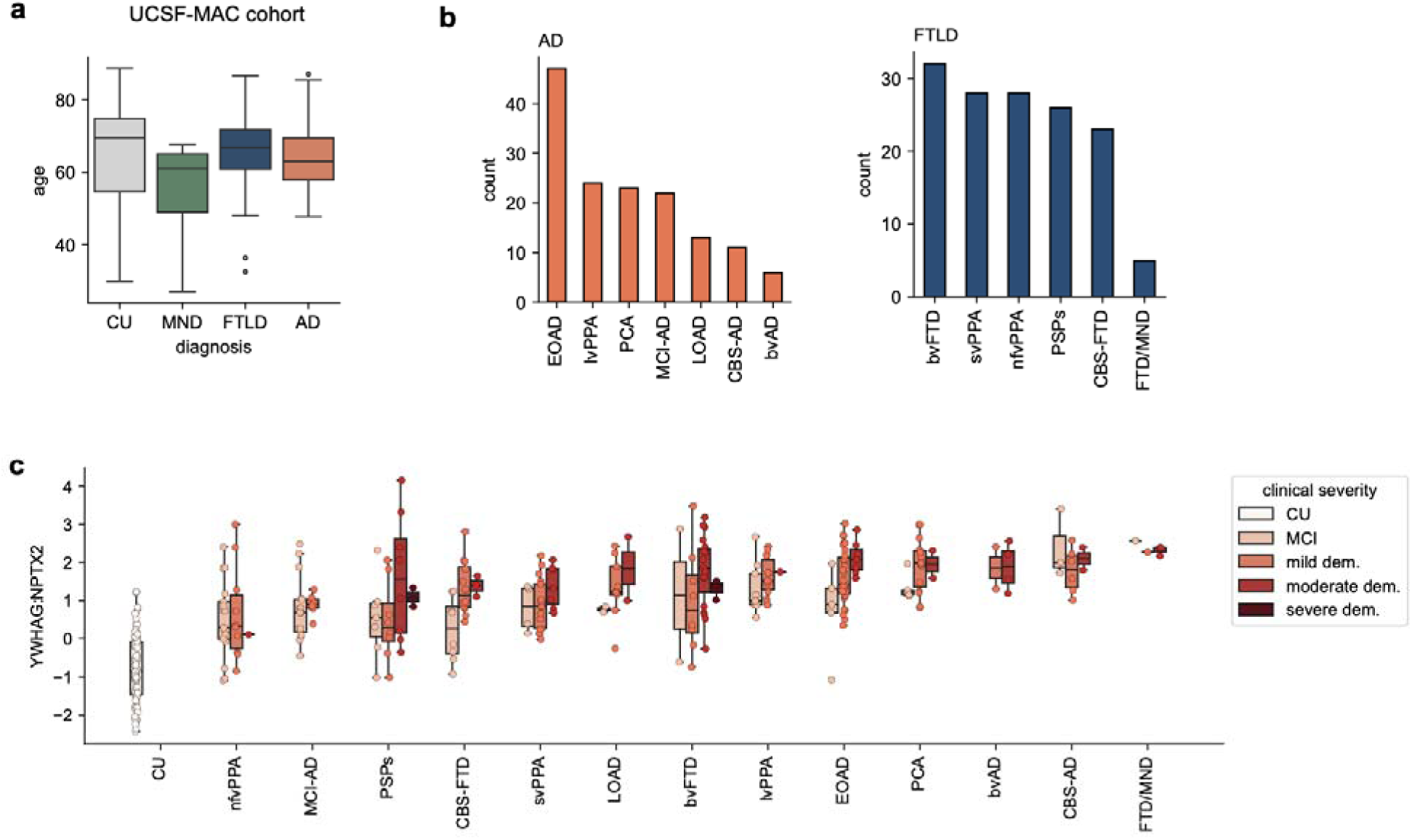
UCSF MAC cohort demographics. **a**, Clinical diagnosis versus age in UCSF-MAC cohort (n=384). **b,** Bar plots showing sample sizes of clinical sub-types of AD (n=144, left) and FTLD (149, right) in the UCSF-MAC cohort. **c,** Levels of CSF YWHAG:NPTX2 in clinical AD and FTLD subtypes, stratified by clinical severity in the UCSF-MAC cohort. For all box plots, box bounds are the Q1, median and Q3; the whiskers show Q1 − 1.5× the interquartile range (IQR) and Q3 + 1.5× the IQR.

**Extended Data Figure 2.**
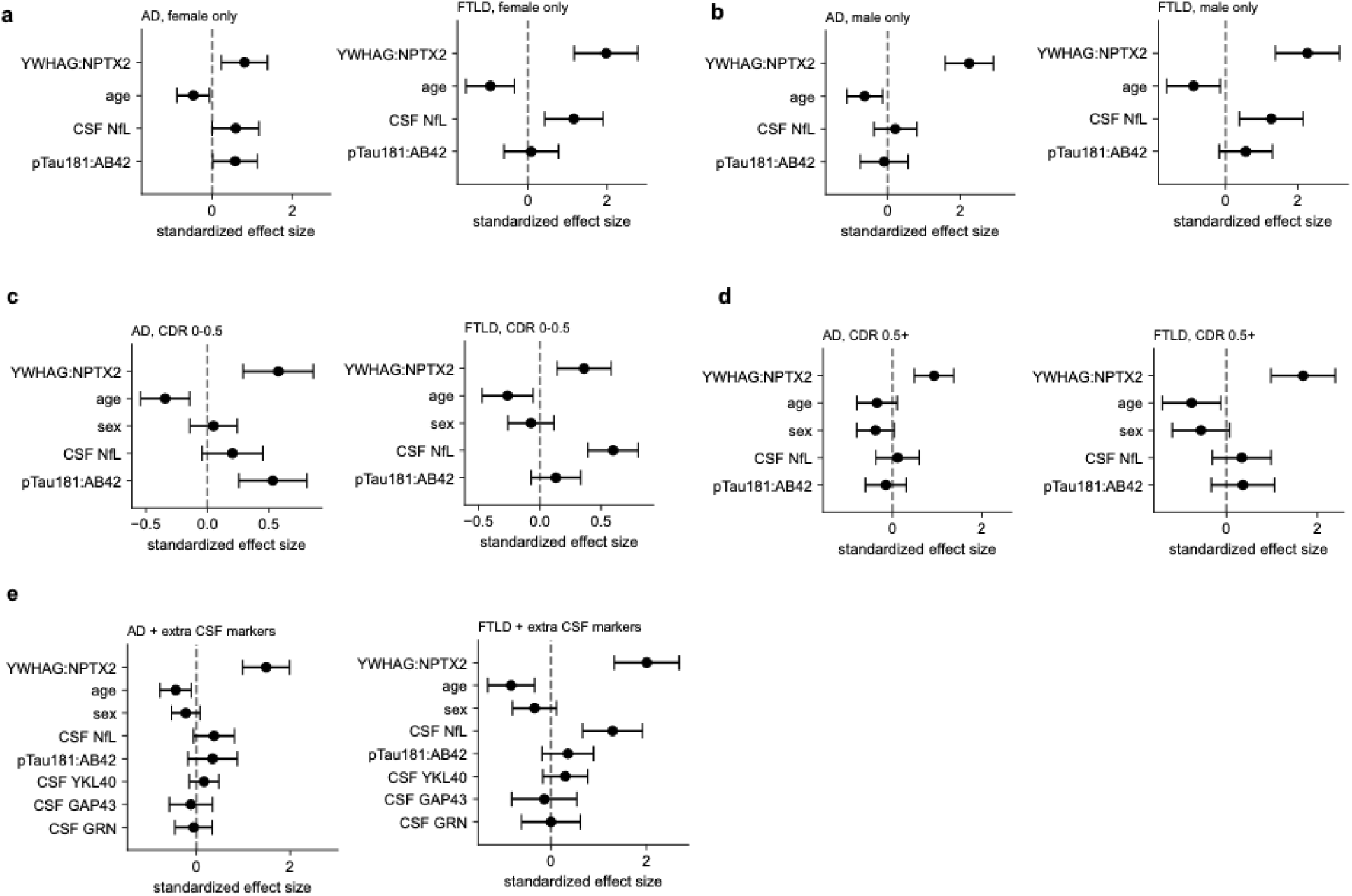
**Sensitivity analyses of the UCSF-MAC clinical severity models.** Each panel shows the AD and control model (left) and the FTLD and control model (right). Points and error bars represent effect sizes and 95% confidence intervals. **a,** Female participants only (AD n=78, FTLD n=51, CU n=23). **b,** Male participants only (AD n=53, FTLD n=48, CU n=23). **c,** CDR Global ≤ 0.5 participants only (AD n=56, FTLD n=73, CU n=46). **d,** CDR Global ≥ 0.5 participants only (AD n=106, FTLD n=99, CU n=0). **e,** All AD-control or FTLD-control participants, but with CSF YKL-40, GAP43, and PGRN included as covariates (AD n=108, FTLD n=145, CU n=46).

**Extended Data Figure 3.**
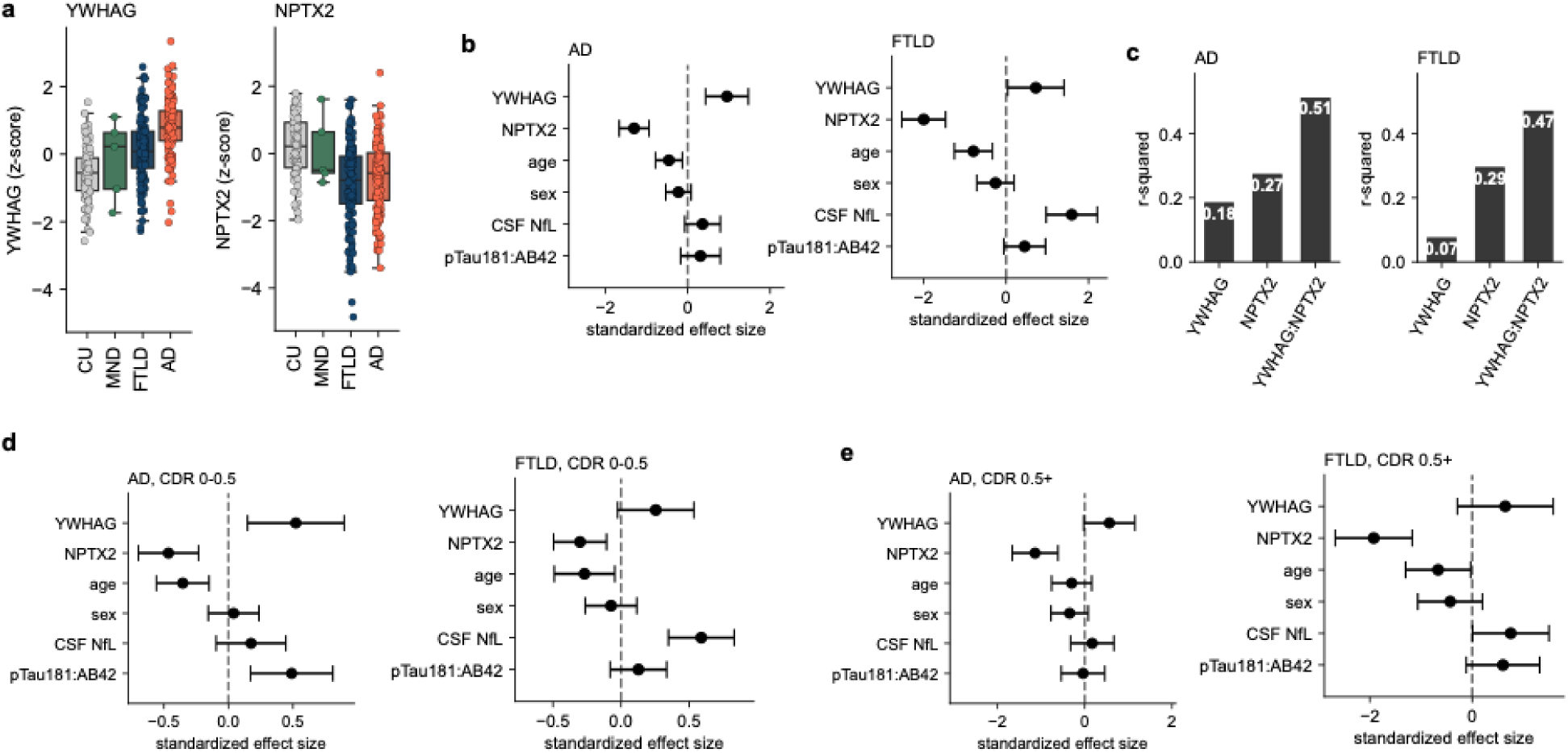
Individual YWHAG and NPTX2 protein analyses in UCSF-MAC. **a**, CSF YWHAG (left) and CSF NPTX2 (right) z-scores by clinical diagnosis in UCSF-MAC (n=381). **b,** CDR Sum of Boxes (AD and control, n=154, left) and FTLD-CDR Sum of Boxes (FTLD and control, n=145, right) regressed against YWHAG and NPTX2 entered as separate predictors alongside age, sex, CSF NfL, and pTau181:Aβ42. **c,** Proportion of variance explained of CDR Sum of Boxes (AD and control, n=154, left) and FTLD-CDR Sum of Boxes (FTLD-control, n=145, right) by CSF YWHAG, NPTX2, or YWHAG:NTPX2. **d,** As in **b**, but restricted to CDR Global ≤ 0.5 participants (AD and control n=102, FTLD n=73). **e,** As in **b**, but restricted to CDR Global ≥ 0.5 participants (AD and control n=106, FTLD n=99).

**Extended Data Figure 4.**
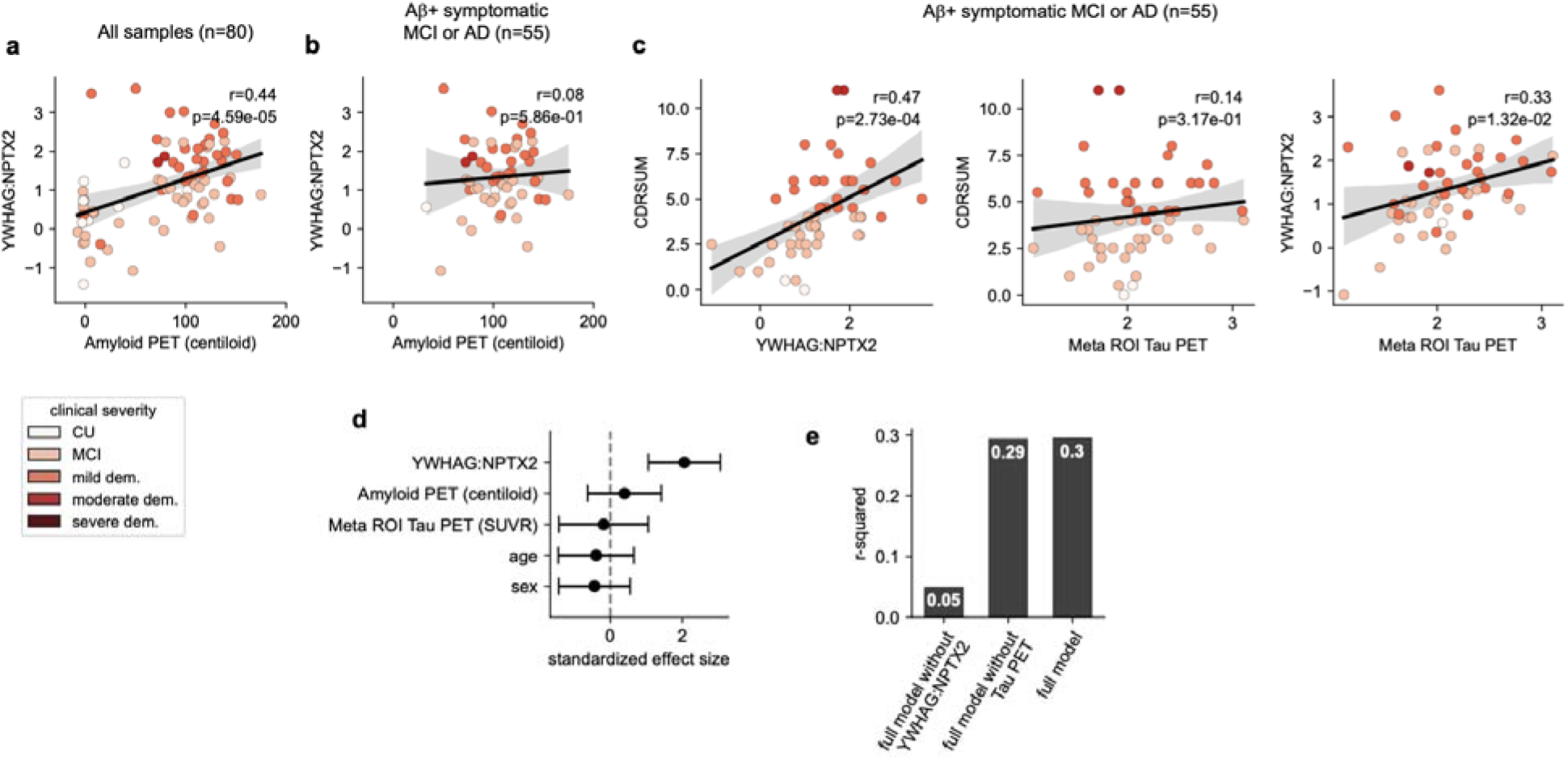
CSF YWHAG:NPTX2 ratio versus amyloid and tau PET in AD. **a**, Linear correlation and p-value between CSF YWHAG:NPTX2 and Amyloid PET (centiloid) in the MAC cohort (n=80), color-coded according to CDR Global. **b,** Linear correlation and p-value between CSF YWHAG:NPTX2 and Amyloid PET (centiloid) in cognitively impaired, amyloid-positive patients in the MAC cohort (n=55), color-coded according to CDR Global. **c,** Pairwise linear correlations and p-values between CDR Sum of Boxes, CSF YWHAG:NPTX2, and tau PET temporal meta-ROI among cognitively impaired, amyloid-positive individuals in the UCSF-MAC cohort (n=55), color-coded according to CDR Global. **d,** CDR Sum of Boxes regressed against CSF YWHAG:NPTX2, age, sex, amyloid PET, and tau PET in a linear model (n=55). Points and error bars represent standardized effect sizes and 95% confidence intervals. **e,** R-squared results from a linear model regressing CDR Sum of Boxes against covariates displayed on the x-axis. Full model includes CSF YWHAG:NPTX2, age, sex, amyloid PET, and tau PET (n=55).

**Extended Data Figure 5.**
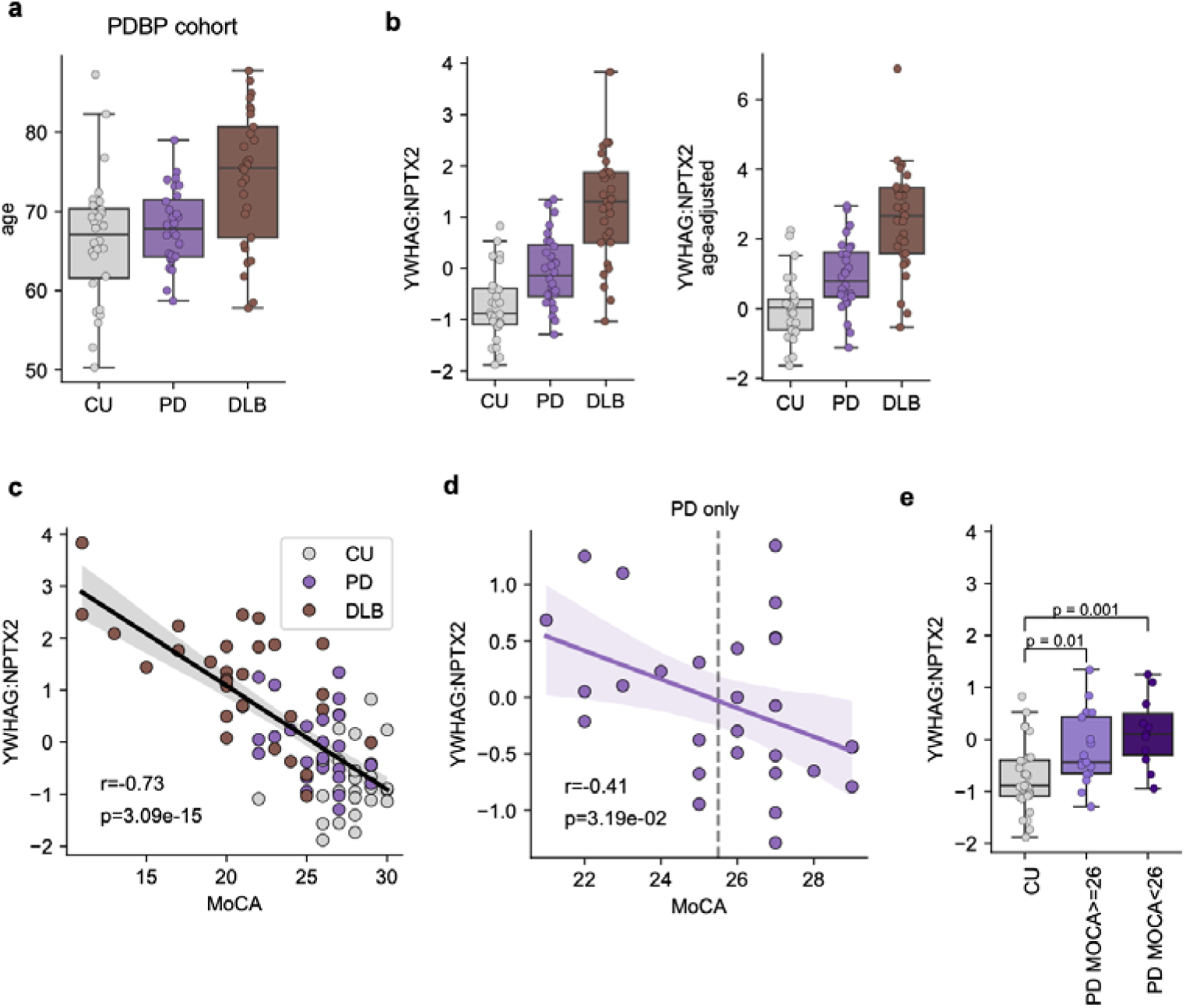
CSF YWHAG:NPTX2 ratio is elevated in Parkinson’s disease and Dementia with Lewy Bodies. **a**, Age per clinical diagnosis in the PDBP cohort (n=85); CU n=28, PD n=28, DLB n=29). **b,** CSF YWHAG:NPTX2 levels (original units left, age-adjusted right) versus clinical diagnosis in PDBP. **c,** Linear correlation and p-value between CSF YWHAG:NPTX2 and MoCA scores (n=85), color-coded according to diagnosis. **d,** Linear correlation and p-value between CSF YWHAG:NPTX2 and MoCA scores in PD subjects (n=28). **e,** CSF YWHAG:NPTX2 levels in PD (n=28) versus control (n=28) subject stratified by MoCA scores. For all box plots, box bounds are the Q1, median and Q3; the whiskers show Q1 − 1.5× the interquartile range (IQR) and Q3 + 1.5× the IQR.

**Extended Data Figure 6.**
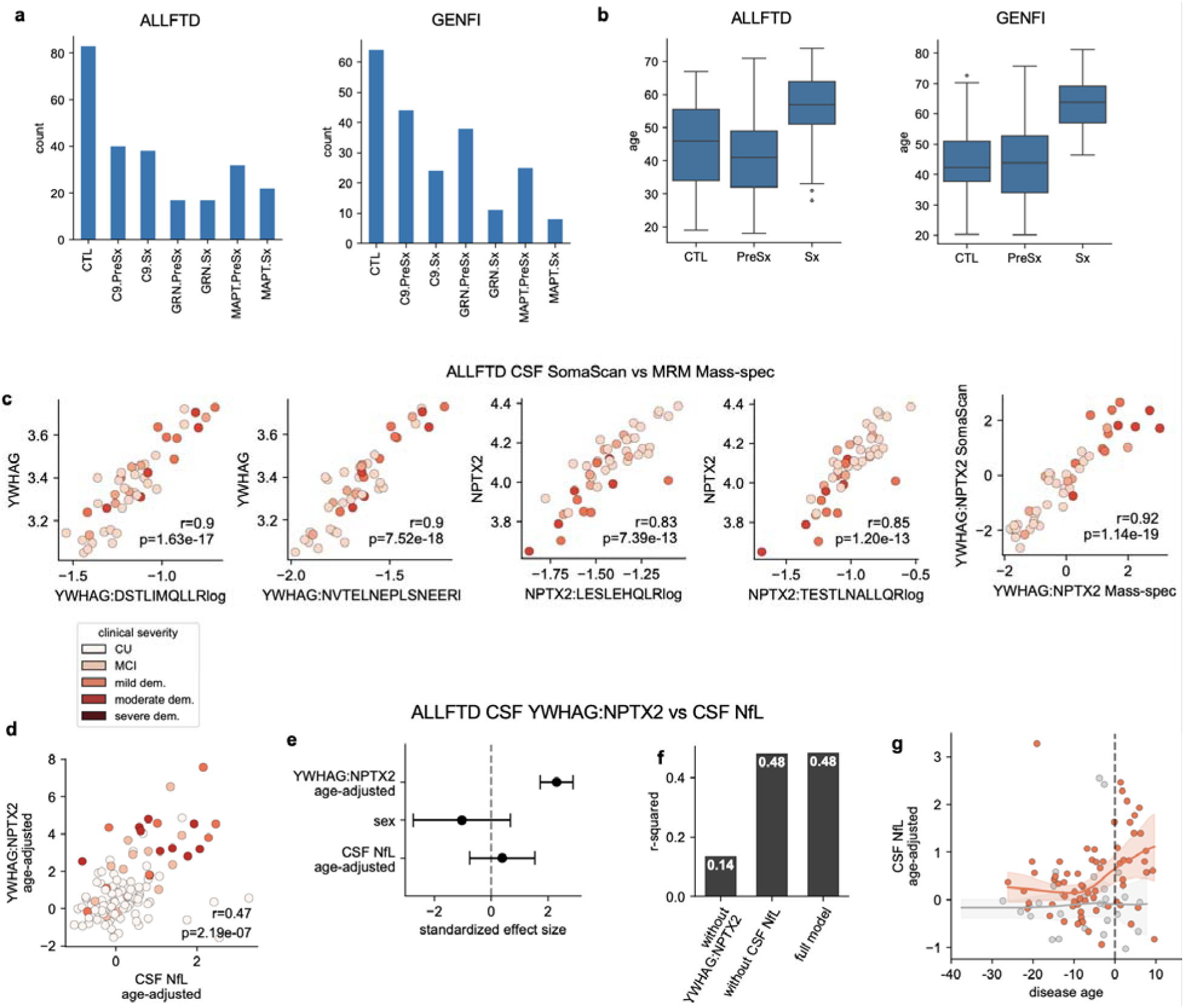
Genetic FTLD cohort demographics and sensitivity analyses. **a**, Sample sizes of familial FTLD gene groups and disease stage in the ALLFTLD (n=249, left) and GENFI (n=214, right) cohorts. **b,** Age of controls, presymptomatic (PreSx), and symptomatic (Sx) genetic FTLD patients in the ALLFTLD (n=249, left) and GENFI (n=214right) cohorts. Box bounds are the Q1, median and Q3; the whiskers show Q1 − 1.5× the interquartile range (IQR) and Q3 + 1.5× the IQR. **c,** Linear correlations and p-values between SomaScan and mass-spectrometry measurements of CSF YWHAG, NPTX2, and YWHAG:NPTX2 in the ALLFTLD cohort, color-coded according to CDR+NACC-FTLD (FTLD-CDR) Global (n=46). Mass-spectrometry CSF YWHAG:NPTX2 represents the ratio between YWHAG:NVTELNEPLSNEER and NPTX2:TESTLNALLQRz. **d,** Linear correlation and p-value between CSF YWHAG:NPTX2 and CSF NfL (Quanterix Simoa, log_10_), color-coded according to CDR+NACC-FTLD (FTLD-CDR) Global (n=109). **e,** FTLD-CDR Sum of Boxes regressed against CSF YWHAG:NPTX2, age, sex, and CSF NfL in a linear model (n=109). Points and error bars represent standardized effect sizes and 95% confidence intervals. **f,** R-squared results from a linear model regressing FTLD-CDR Sum of Boxes against covariates displayed on the x-axis (n=109). Full model includes CSF YWHAG:NPTX2, age, sex, and CSF NfL. **g,** Changes with estimated years until FTLD symptom onset of age-adjusted CSF NfL (n=97; 29 non-carriers, 68 carriers). Lowess regression lines with 95% confidence interval for mutation carriers (orange) versus non-carriers (blue) are shown.

**Extended Data Figure 7.**
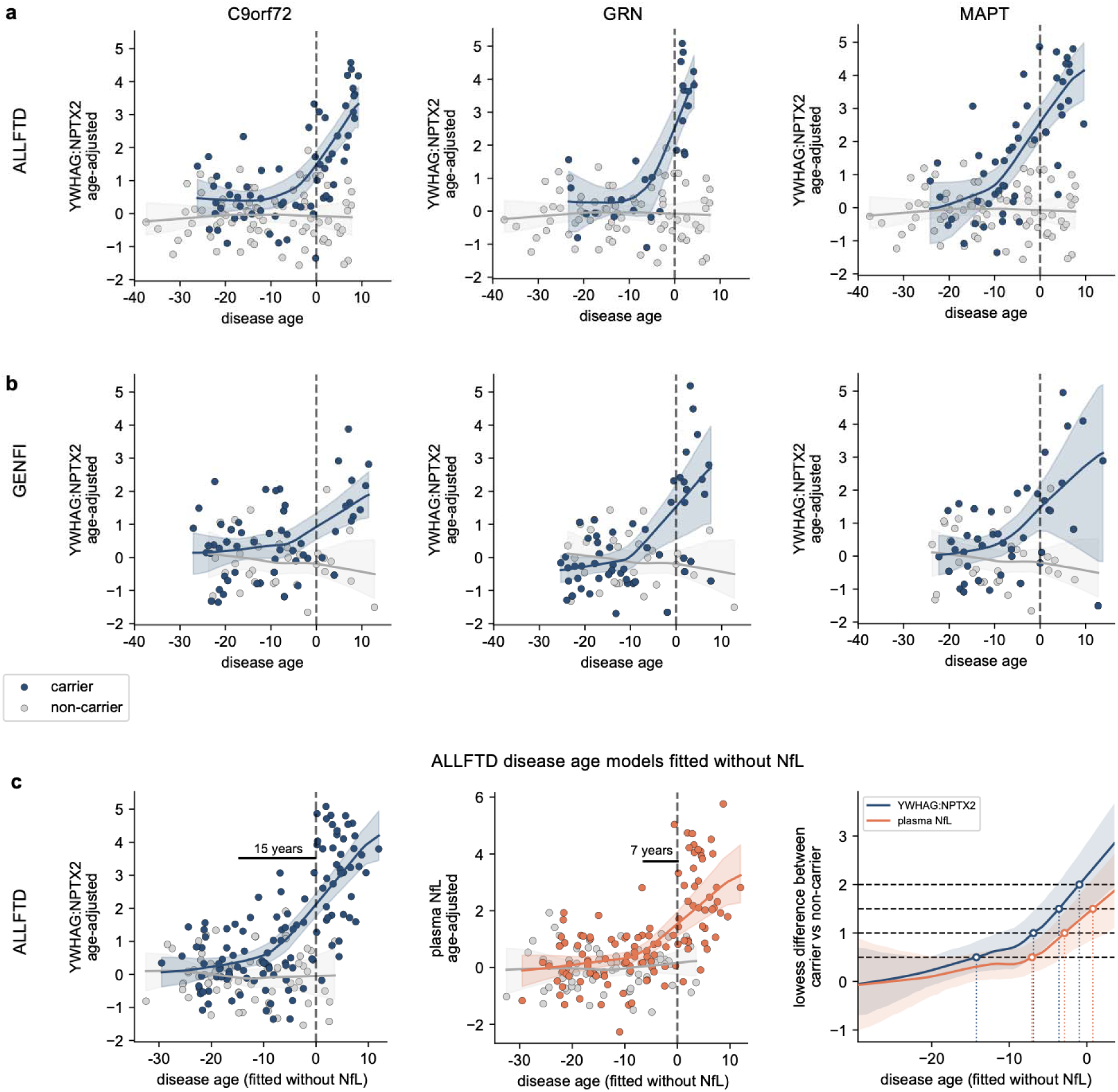
CSF YWHAG:NPTX2 ratio versus estimated years until symptom onset per genetic FTLD gene group and a disease age axis fitted without NfL. **a**, Changes with estimated years until FTLD symptom onset of CSF YWHAG:NPTX2, stratified by mutation-carrier status and gene group (C9orf72 n=63 left, GRN n=28 middle, MAPT n=49 right against non-carriers n=77) in the ALLFTD cohort. Lowess regression lines with 95% confidence interval for carriers (orange) versus non-carriers (blue) are shown. **b,** As in **a,** but in the GENFI cohort (C9orf72 n=55 left, GRN n=53 middle, MAPT n=42 right against non-carriers n=44). **c**, The ALLFTD Fig. 3a**-c** analyses repeated on a disease-age axis refitted without plasma NfL among its inputs (n=188).

**Extended Data Figure 8.**
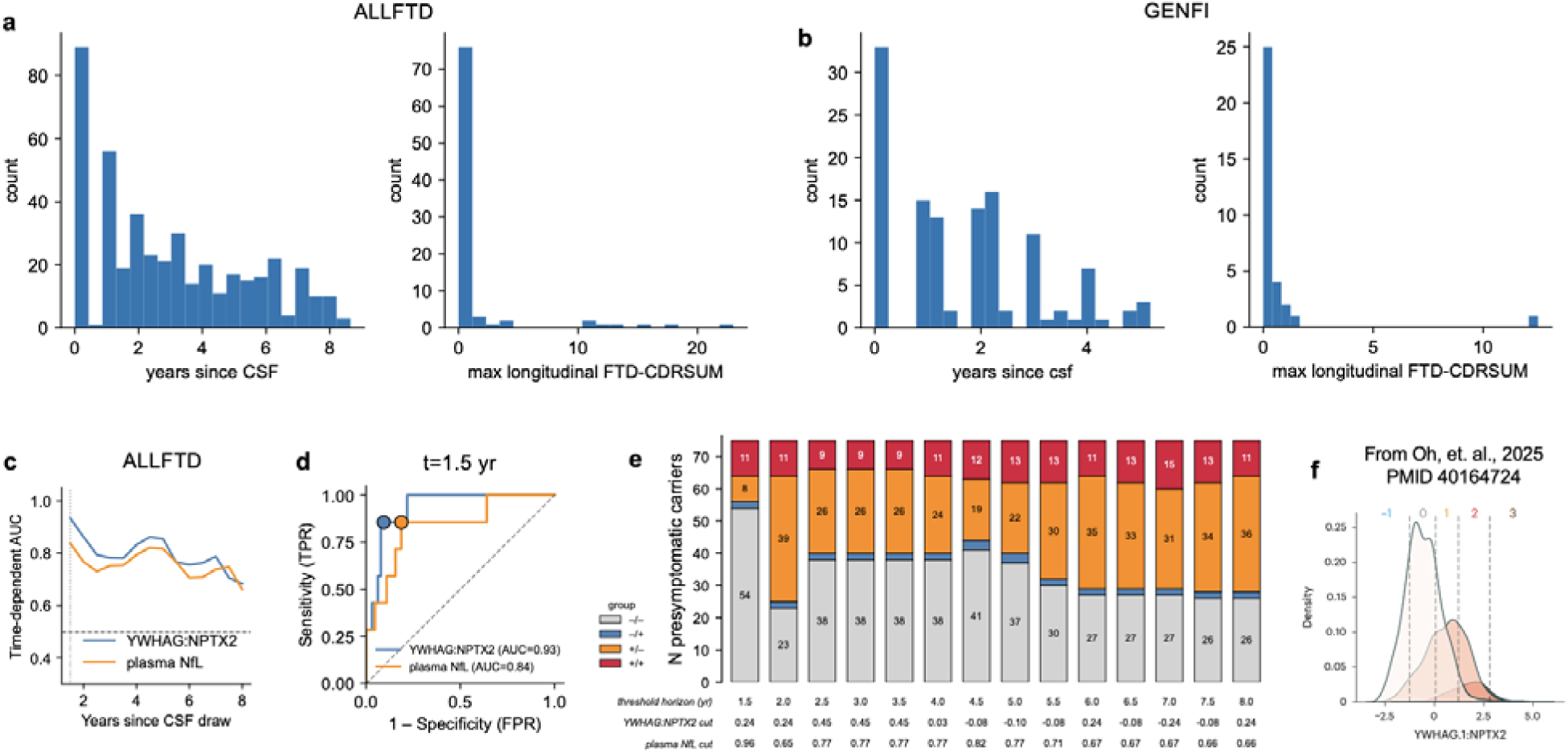
Longitudinal cognitive follow-up and derivation of biomarker thresholds. **a**, Histogram showing distribution of follow-up years (left) and maximum CDR+NACC-FTLD (FTLD-CDR) Sum of Boxes over all years among presymptomatic carriers in the ALLFTLD cohort (n=75). **b,** As in **a**, but for the GENFI cohort (n=46). **c,** Time-dependent ROC-AUC for CSF YWHAG:NPTX2 and plasma NfL predicting phenoconversion, across follow-up horizons from 1.5 to 8 years (n=75). Both markers peak at 1.5 years (AUC 0.93 and 0.84). **d,** ROC curves at the 1.5-year horizon with the selected operating points marked: plasma NfL at >80% sensitivity and CSF YWHAG:NPTX2 at >90% specificity. **e,** Number of presymptomatic carriers falling into each biomarker group quadrant when both thresholds are re-derived at each follow-up horizon. The horizon and two biomarker thresholds are noted beneath each bar. **f,** AD disease stage thresholds for CSF YWHAG:NPTX2 derived from Oh et. al., *Nature Medicine*, 2025.

**Extended Data Figure 9.**
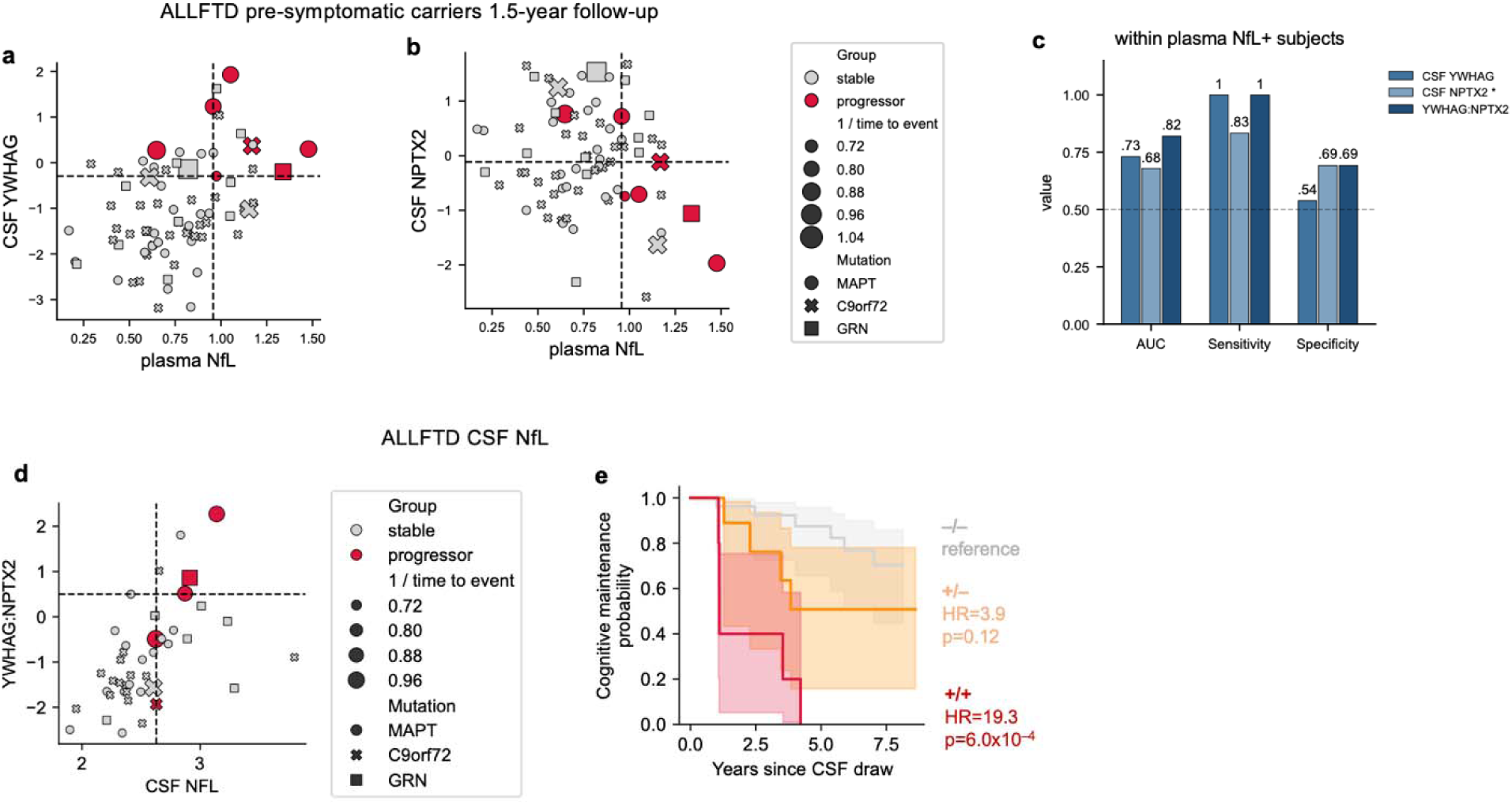
CSF YWHAG:NPTX2 versus individual proteins and CSF NfL in predicting presymptomatic phenoconversion. **a**, CSF YWHAG versus plasma NfL in ALLFTD presymptomatic carriers, with thresholds derived as done with CSF YWHAG:NPTX2 and points color-coded by phenoconversion within 1.5 years (n=75). **b,** As in **a,** but for CSF NPTX2. **c,** Discrimination of 1.5-year phenoconverters within plasma NfL-positive carriers by CSF YWHAG:NPTX2, YWHAG alone, or NPTX2 alone. **d,** Scatterplot showing CSF YWHAG:NPTX2 versus CSF NfL in ALLFTLD presymptomatic carriers, with points color-coded by phenoconversion within 1.5 years, sized by time until conversion, and shaped by mutation (n=42). Dotted lines define positive and negative biomarker gates determined as in **Extended Data** Fig. 8d. **e,** Kaplan Meier curves with 95% confidence intervals showing symptom onset over 8 follow-up years in presymptomatic carriers in the ALLFTLD cohort, stratified by biomarker groups defined by gates in **d**. Hazard ratios and p-values from a Cox proportional hazard model controlling for age and sex are shown.

**Extended Data Figure 10.**
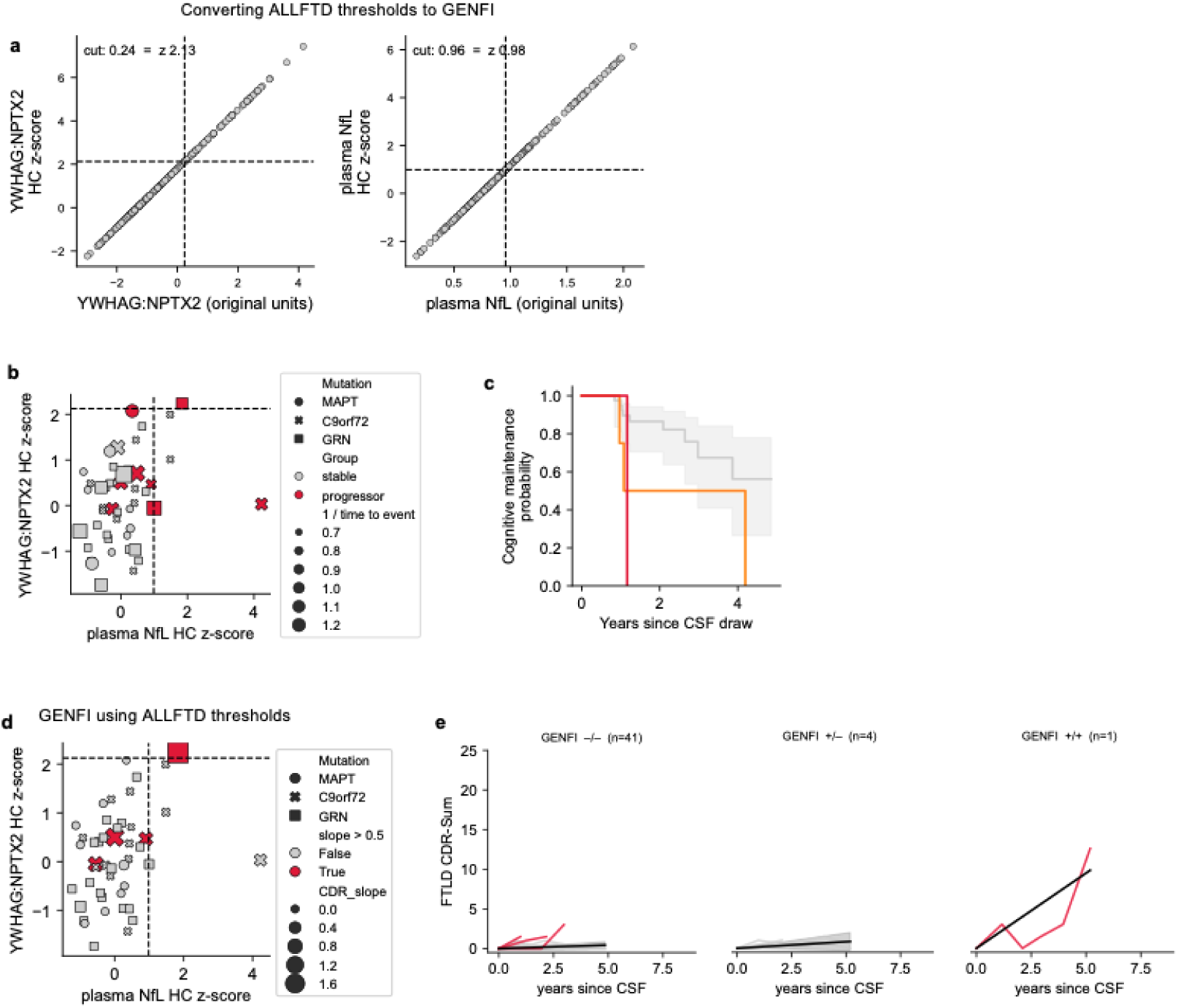
Applying ALLFTD-derived thresholds to GENFI. **a**, Conversion of the ALLFTD thresholds for CSF YWHAG:NPTX2 (left) and plasma NfL (right) onto the healthy-control z-scale, shown in z versus original units. **b,** CSF YWHAG:NPTX2 versus plasma NfL in GENFI presymptomatic carriers (n=46) with points color-coded by phenoconversion within 1.5 years, sized by time until conversion, shaped by mutation, and the transferred thresholds shown. **c,** Kaplan-Meier curves for phenoconversion by groups based on **b**. **d,** As in **b**, but with points color-coded and sized by by longitudinal slope of FLTD CDR-Sum of Boxes. **e,** Spaghetti plot showing longitudinal trajectories in CDR+NACC-FTLD (FTLD CDR) Sum of Boxes stratified by biomarker group over 8 follow-up years in GENFI. Average slopes with 95% confidence intervals are shown.

**Supplementary Figure 1.**
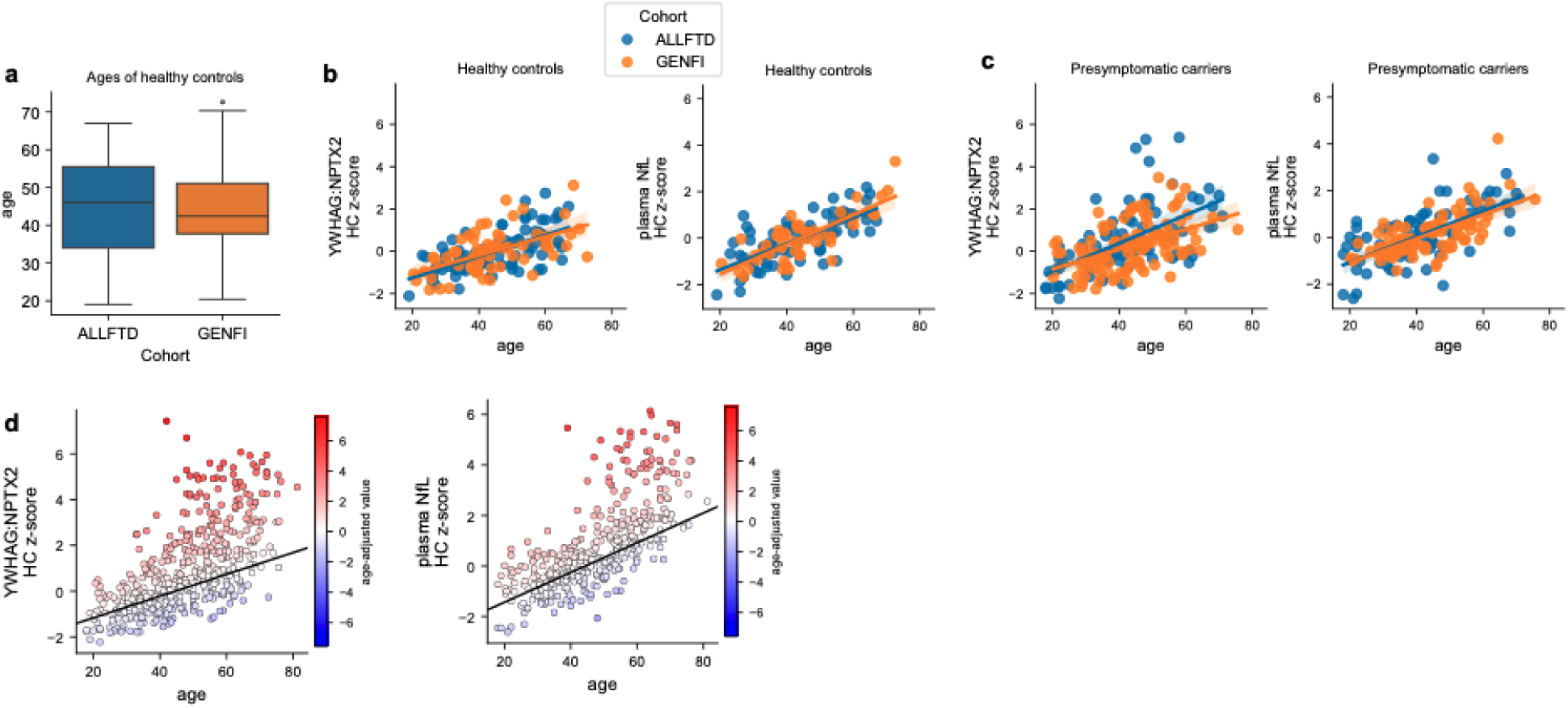
Cross-cohort harmonization and normalization of CSF YWHAG:NTPX2 and plasma NfL in ALLFTD and GENFI. **a**, Age of healthy controls in ALLFTD (n=83) and GENFI (n=64). The box bounds are the Q1, median and Q3; the whiskers show Q1 − 1.5× the interquartile range (IQR) and Q3 + 1.5× the IQR. **b,** Healthy control z-scored CSF YWHAG:NPTX2 (left) and plasma NfL (right) versus age in healthy controls from both cohorts, with cohort-specific regression lines and 95% confidence intervals. **c,** As in **b**, but with presymptomatic carriers (ALLFTD n=89, GENFI n=107). **d,** All participants from both cohorts (ALLFTD n=249, GENFI n=214), with points color-coded by the age-adjusted value of each marker – the residual from that participant’s own cohort’s healthy control age regression. The black line is the pooled healthy control age regression line.

**Supplementary Figure 2.**
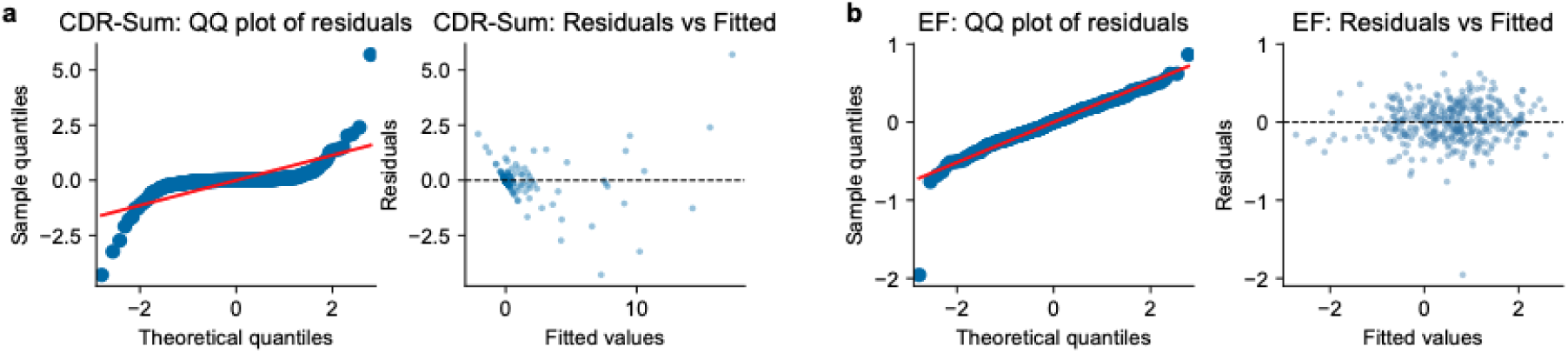
Mixed-effects model residual diagnostics in ALLFTD. **a**, Residual diagnostics for the FTLD-CDR Sum of Boxes model in Fig. 4d (n=73). **b,** Residual diagnostics for the UDS3-EF model in Fig. 4g (n=73).

**Supplementary Figure 3.**
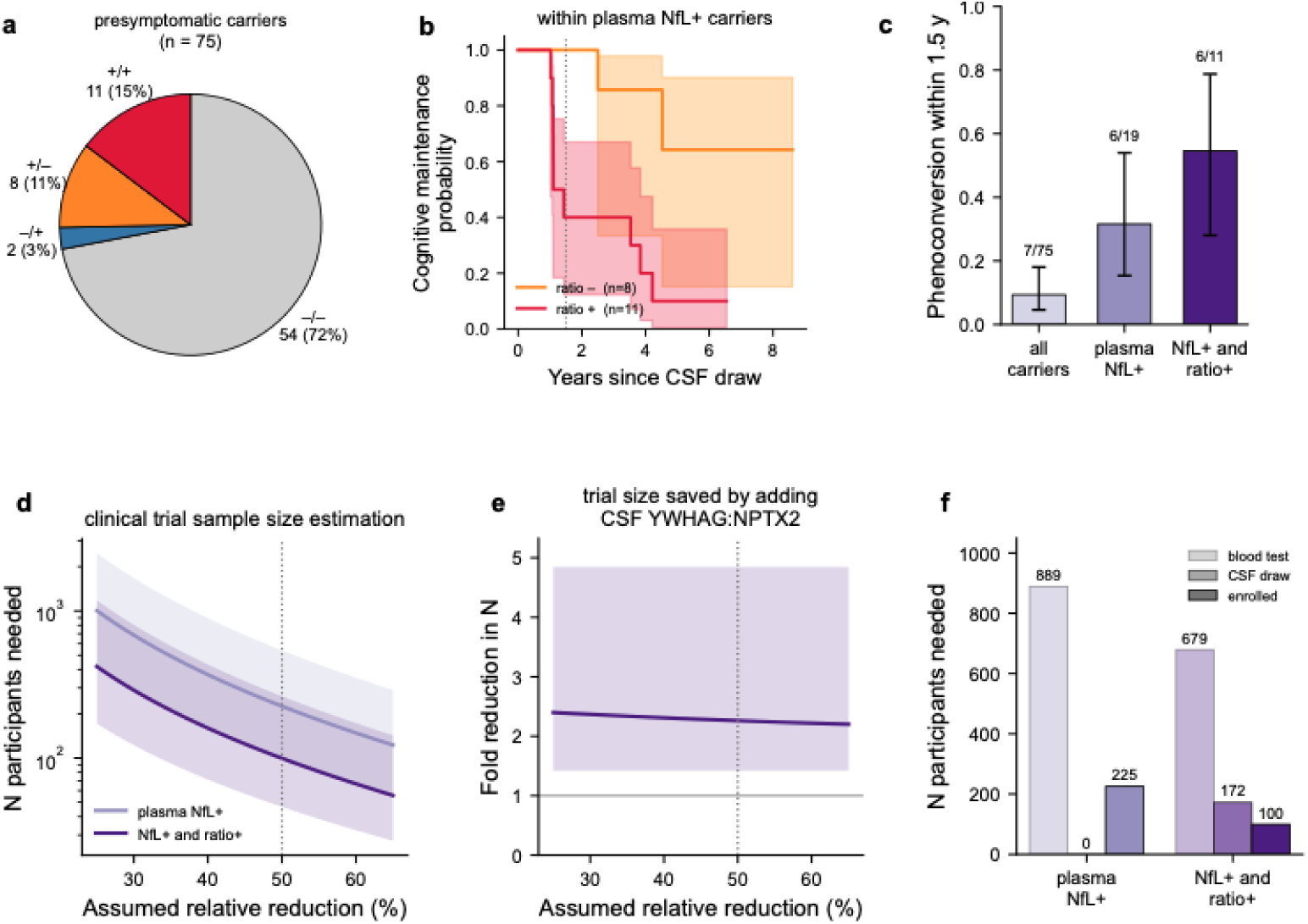
Projected effect of two-tiered biomarker screening on FTLD prevention trial enrollment. **a**, Distribution of the 75 ALLFTD presymptomatic carriers across the four biomarker groups in Fig. 4a. **b,** Kaplan-Meier curves for phenoconversion within plasma NfL-positive carriers, stratified by CSF YWHAG:NPTX2 positivity. **c,** Observed 1.5-year phenoconversion rate under each screening strategy, with 95% Wilson score intervals. **d,** Total participants required across both arms as a function of the assumed relative reduction in phenoconversion, for a two-arm 1:1 trial at two-sided α = 0.05 and 80% power. Bands span the required N implied by the 95% interval of the observed event rate. **e**, Fold reduction in required sample size from adding CSF YWHAG:NPTX2 to a plasma NfL screen, with a 95% confidence interval. **f**, Blood tests, CSF draws, and enrollments required under each strategy.

